# IgG autoantibodies in bullous pemphigoid directly induce a pathogenic MyD88-dependent pro-inflammatory response in keratinocytes

**DOI:** 10.1101/2024.10.07.616103

**Authors:** Lei Bao, Christian F. Guerrero Juarez, Jing Li, Manuela Pigors, Shirin Emtenani, Yingzi Liu, Aadil Ahmed, N Ishii, T Hashimoto, Bethany E. Perez White, Stefan Green, Kevin Kunstman, Nicole C Nowak, Connor Cole, Virgilia Macias, Maria Sverdlov, M. Allen McAlexander, Christopher McCrae, Christopher D. Nazaroff, Enno Schmidt, Kyle T. Amber

## Abstract

While autoantibodies in bullous pemphigoid (BP) are known to activate the innate immune response, their direct effect on keratinocytes, and the contribution of BP-IgG autoantibody-dependent keratinocyte responses to BP pathology is largely unknown. Herein, we performed multiplex immunoassays and bulk RNA-seq on primary keratinocytes treated with IgG from BP patients or controls. We identified a pro-inflammatory and proteolytic response with release of several cytokines (IL-6, IL-24, TGF-β1), chemokines (CXCL16, CTACK, MIP-3β, RANTES), C1s, DPP4, and MMP-9. We further validated this response using spatial transcriptomics and scRNA-seq of diseased and control skin. Blistering itself appeared to be major driver of this inflammatory response, with attached BP skin and spongiotic dermatitis revealing highly similar transcriptomes. Based on elevated levels of MyD88 and MyD88-dependent cytokines, we studied the impact of MyD88 deficiency in keratinocytes and demonstrated that MyD88 regulates BP-IgG-induced expression of IL-8, IL-24, and MMP-9. Induction of experimental BP in mice with *Krt14*-specific *Myd88* knockout revealed significantly decreased disease severity with decreased serum levels of IL-1β, IL-4, and IL-9 indicating the contributory role of keratinocyte-derived skin inflammation towards systemic response. Our work demonstrates the key contributions of keratinocyte and MyD88 dependent signaling in response to autoantibodies in BP.

**Key Messages:** -IgG antibodies from bullous pemphigoid (BP) patients induce significant upregulation of several inflammatory markers in keratinocytes including cytokines (IL-6, IL-24, TGF-β1), chemokines (CXCL16, CTACK, MIP-3β, RANTES), C1s, DPP4, and MMP9. Several of these markers, including IL-8, IL-24, and MMP9 are regulated by MyD88.

-Spatial transcriptomics reveals that BP patient blistered skin demonstrated similar transcriptomic profiles to BP-IgG-treated keratinocytes. With attached skin demonstrating a comparable transcriptome to that seen in spongiotic dermatitis.

-In a mouse BP model, keratinocyte-specific MyD88 deficiency results in decreased disease severity with a subsequent decrease in serum IL-1β, IL-4, and IL-9 levels.

**Capsule summary:** IgG from patients with bullous pemphigoid (BP) induces a pro-inflammatory response in keratinocytes, indicating their direct role in driving the inflammatory response in BP.

## Introduction

Bullous pemphigoid (BP) is an autoimmune blistering disease caused by autoantibodies targeting the hemidesmosomal proteins BP180 (type 17 collagen, COL17) and to a lesser extent, BP230^1, 2^. Autoantibodies can be internalized in a BP-IgG/BP180 complex via macropinocytosis^3, 4^, as well as interact with Fc receptors (FcR) and complement^5–7^. Clinically, BP presents with numerous morphologies including urticarial plaques, eczematous patches, and/or bullae, with accompanying pruritus^1^. Histologically, BP most commonly presents with abundant tissue eosinophilia, particularly with eosinophils lining the dermo-epidermal junction in early (urticarial) phase and/or entering the inflamed epidermis (eosinophilic spongiosis)^8, 9^. Both complement-fixing and non-fixing IgG isotypes are seen amongst autoantibodies^5, 6, 10, 11^. Aside from IgG autoantibodies, IgE autoantibodies are detected in up to 70% of patients, driving basophil degranulation^5, 8, 12–14^. Both IgG and IgE from patients with BP have been shown to cause the release of IL-6 and IL-8 directly from cultured keratinocytes, indicating an FcR and complement-independent pathologic mechanism^15–17^. This suggests that autoantibody-triggered manipulation of BP180 on the keratinocyte may regulate inflammatory pathways, and directly initiate and/or amplify the local inflammatory response in BP. IL-6 and IL-8 upregulation alone would, however, not sufficiently explain the lesional Th2 polarization^18, 19^ and eosinophil influx seen in BP. Thus, we hypothesize that IgG from BP patients activates alternative pathologic pathways. For example, we have recently discovered direct inflammatory effects and pauci-inflammatory blistering mechanisms of autoantibodies in anti-laminin 332 pemphigoid^20, 21^.

Despite the known roles of BP180 in the basement membrane zone (BMZ), as an autoantigen in BP, and as a prognostic marker in certain cancers^22^, its role in the regulation of skin inflammation and BMZ homeostasis is not well described^23^. Several animal mutant models of *Col17a1*^24, 25^, as well as case reports of epidermolysis bullosa with *COL17A1* mutations^26–28^ strongly suggest a direct regulatory role of COL17 on tissue granulocyte infiltration. COL17 additionally undergoes physiological ectodomain shedding^29^, which has been associated with carcinogenesis^30^, but has an unclear role in skin inflammation^22, 23, 31–34^. Notably, spontaneous inflammation occurs in two animal models with a mutation placed in the humanized COL17 NC16a domain^24^, or the corresponding mouse NC14a domain^25^. In the first model, mice developed spontaneous scratching behavior even when crossed with *Rag2* -/- mice^24^, indicating a lack of adaptive immune involvement. Cultured keratinocytes from these ΔNC16a mice demonstrated elevated TSLP relative to controls. The study also noted a significant increase in TSLP in BP skin relative to normal, indicating TSLP as an epidermal contributor towards pathogenesis. We, however, noted epithelial TSLP elevation to be a non-specific finding of multiple inflammatory skin diseases^35^. In the second disease model, introducing a mutation in NC14a (ΔNC14A)^25^ also resulted in a BP-like phenotype with dermal eosinophil infiltrations, peripheral eosinophilia, elevated serum IgE, and pruritus^25^. Oddly, despite the removal of the major BP epitope region, some ΔNC14A mice developed IgG and IgA autoantibodies with subepidermal reactivity. These IgG autoantibodies recognized a 180-kDa keratinocyte protein that was sensitive to collagenase digestion, consistent with COL17. Thus, it is unclear to what extent the symptomatology is due to direct *COL17* mutations or the generation of anti-skin antibodies in this model.

In light of these studies, we hypothesized that disturbance of BP180 homeostasis by autoantibodies in BP drives an inflammatory response from keratinocytes, extending beyond IL-6 and IL-8. In this study, we investigated the effects of IgG from patients with BP on human keratinocytes *in vitro* using a transcriptomic approach complemented with large-scale validation of inflammatory molecules at the protein level. We demonstrated that many of these inflammatory changes depend on the Myeloid Differentiation primary response 88 (MyD88) protein. We relate this keratinocyte-dependent inflammatory response to patient pathology by correlating the *in vitro* response to spatial transcriptomic and scRNA-seq data of epithelia from patients with BP. Further, we demonstrate the *in vivo* relevance of this response using a recently described mouse model of BP^36^ where we show that keratinocyte-specific knockout of MyD88 was sufficient to mitigate disease severity and reduce circulating type 2 inflammatory cytokines such as IL-4 and IL-9.

## Materials and Methods

### Ethics approvals

Studies utilizing human samples were approved by the Institutional Review Board at Rush University Medical Center (#20121406) and conducted according to the principle of the Declaration of Helsinki. Animal studies were approved by the Institutional Animal Care and Use Committee at Rush University Medical Center (#20-079, # 22-024) in accordance with NIH guidelines.

### Patients

Blood was collected from patients with active BP following written informed consent. A diagnosis of BP was confirmed using previously reported standards^37^ including clinical suspicion of BP, positive direct immunofluorescence, and a positive serologic test either for BP180 and/or BP230 autoantibodies by ELISA, and/or deposition on the roof of a salt split skin blister.

### Mice

B6N.Cg-Tg (*Krt14*-cre)1Amc/J mice were crossed for at least 6 generations with C57B6/J mice (referred to in the text as *Krt14*-Cre). *Myd88^fl/fl^ (*B6.129P2(SJL)-*Myd88*tm1Defr/J) mice were generated as previously described^38^. This strain contains a loxP site flanking exon 3 of *Myd88* which retains MyD88 function in the absence of Cre recombinase. When homozygous mice are bred to Cre-expressing mice, they demonstrate deletion of exon 3 in *Myd88*. All strains (JAX stock No.018964, 000664 and 008888, respectively) were purchased from Jackson Laboratories (Bar Harbor, Maine, USA). Genotyping was performed by TransnetYX. Animals were housed under specific pathogen-free conditions with a 12-h light–dark cycle and fed standard chow *ad libitum*.

### Multiplex Immunoassay and ELISA

Multiplex measurements of supernatants were performed using the Luminex 200 system (Luminex, Austin, TX, USA) by Eve Technologies Corp. (Calgary, Alberta, Canada). TGF-β 3-Plex Discovery Assay® (MilliporeSigma, Burlington, Massachusetts, USA), Eve Technologies’ Human MMP/TIMP 13-Plex Discovery Assay, and Eve Technologies’ Human Cytokine 96-Plex Discovery Assay were used to measure supernatant cytokine, chemokine, and metalloprotease expression according to the manufacturer’s instructions. Additional specific markers including ADAM8 (Biorbyt, Durham, North Carolina, USA), IL-36G (Thermo Fisher Scientific, Waltham, Massachusetts, USA), C1s (RayBiotech, Peachtree, Corners, Georgia, USA), and CD26 (Thermo Fisher Scientific) were measured by ELISA following the manufacturer’s instructions. Mouse cytokines were measured using the 32-plex cytokine assay (MilliporeSigma). Standard curves were generated for each analyte to define the concentrations of each analyte. All samples were run in duplicate, with the average of the duplicates used for further analysis. A complete list of analytes assessed is shown in the supplementary materials and methods.

### 2D Cell Culture and Treatments

Serum underwent affinity purification using ToxOut (BioVision, Milpitas, California, USA) or Nab columns (Thermo Fisher Scientific), followed by buffer exchange and concentration using Amicon Ultra-15 centrifugal filter units with a 50kDa filter in PBS (Millipore Sigma,) with magnetic azide removal (Nanopartz, Loveland, CO). Multiple purchased pooled human control IgG and individual normal serum controls were utilized to increase biological variability of control IgG. Pooled IgG was purchased from MP Biomedicals (Santa Ana, California, USA), Cell Sciences (Newburyport, Massachusetts, USA), and Sigma (Saint Louis, Missouri, USA). Individual donor sera were purchased from Innovative Research (Novi, Michigan, USA) and underwent the same purification as BP samples. Primary adult human keratinocytes (PHKs) (Thermo Fisher Scientific) were cultured in EpiLife Medium with Human Keratinocyte Growth Supplement (both from Thermo Fisher Scientific) in a humidified atmosphere of 5% CO2 at 37°C. MyD88 knockout N/TERT cells or CRISPR/Cas9 transfection controls were generated as previously described, ^39^, and kindly provided by Dr. Gudjonsson, with the permission of Dr. James G. Rheinwald^40^. N/TERT cells were grown in KC-SFM medium (Thermo Fisher Scientific) supplemented with 30 μg/ml bovine pituitary extract, 0.2 ng/ml epidermal growth factor, and 0.3 mM calcium chloride. When confluency reached approximately 70%, cells were treated with patient or control-IgG at a final concentration of 4.0 μg/μl overnight. Each sample represents a distinct culture. Supernatants were removed and stored at −80°C until protein measurement. Cells pellets were washed twice with PBS and stored at −80°C until RNA extraction. Independent experiments were performed on different passages.

### 3D Human Skin Equivalents

3D human skin equivalents (HSE) were generated and propagated as previously described using primary neonatal human keratinocytes^41^. Three unique donors were used for replicate 3D HSE. After 9 days of growth at the air-liquid interface, 3D HSE were either treated with 2mg/mL of pooled BP-IgG from 4 donors with BP180 reactivity or pooled control-IgG for 72 hours. All cultures were harvested on day 12. Sections were either fixed in formalin or placed in OCT compound with routine histology, or direct immunofluorescence performed by standard procedures.

Detailed methods on RNA extraction, RT-PCR, bulk RNA-seq, tissue microarray generation, spatial transcriptomics, and bioinformatics are provided in the supplemental methods.

## Results

### Pro-inflammatory Cytokines and Chemokines are Released from BP-IgG treated Primary Human Keratinocytes

To investigate the direct effect of BP-IgG on keratinocytes, we cultured primary human keratinocytes (PHK) overnight with affinity purified single donor BP-IgG or control-IgG. The demographics and serotypes of the patients are summarized in (**Supplemental Table 1**). We first analyzed protein levels of 118 cytokines, chemokines, and proteases from keratinocyte supernatants from two independent experiments (**Figure 1A**). Analytes falling below the linear detection range were excluded from further analysis (**Supplemental Data 1**). Setting a threshold with correction for false discovery rates of P_adj_ < 0.05, revealed upregulation of several cytokines (IL-6, IL-24, TGF-β1), chemokines (CXCL16, CTACK, MIP-3β, RANTES), complement components (C1s), DPP4, and proteases (MMP-9) (**Figure 1B**). To better understand co-expression, we next generated a correlation matrix which highlighted a strong correlation between CTACK, IL-8, IL-6, I-TAC, IL-1α, EN-78 and MIP-3β expression (**Figure 1C**). Subanalysis of PHK treated with IgG from patients with only anti-BP180 or anti-BP230 IgG, but not both, revealed comparable findings, other than reduced G-CSF, IL-8, MCP-1, and TNF-α in anti-BP230 IgG treated keratinocytes (**Figure 1D**). Given the elevated MMP-9 production and known role of anti-BP180 IgG in inducing BP180 internalization, we sought to confirm the ability of BP-IgG to induce blistering in 3D HSE in the absence of immune cells using human 3D skin equivalents. This revealed that BP-IgG alone was sufficient to induce blistering in the absence of immune cells (Figure 1E, F).

**Figure 1:**
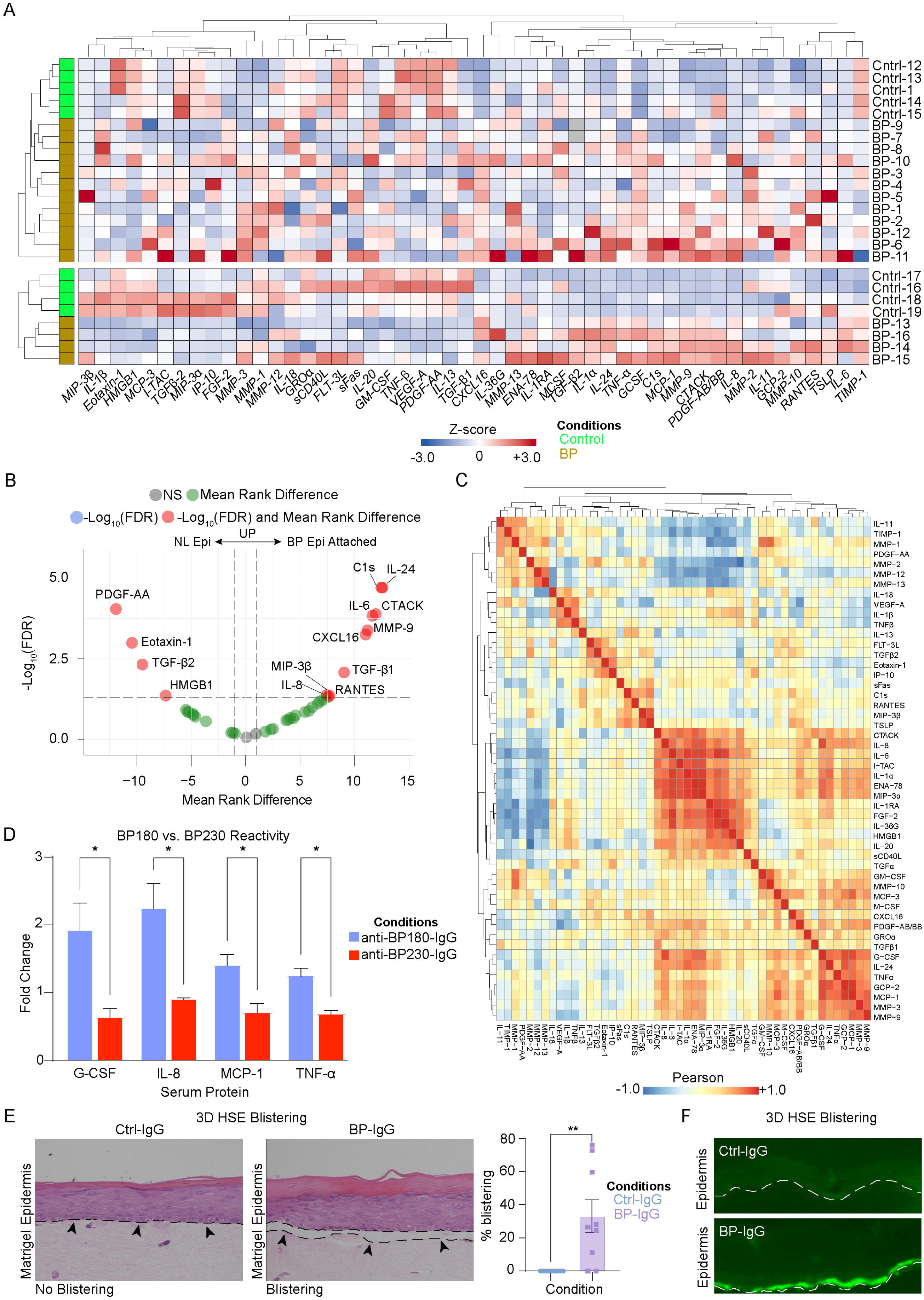
BP-IgG induces expression of numerous pro-inflammatory proteins relative to control-IgG treated primary human keratinocytes. **A**) Heat map demonstrating supernatant protein expression of BP-IgG vs. control-IgG-treated primary human keratinocytes. **B)** Volcano plot of significant up-and down-regulated proteins with *P*_adj_ < 0.05. Mann-Whitney U-test with False Discovery Rate (FDR) correction with Benjamini, Krieger, and Yekutieli method**. C)** Correlation matrix of supernatant protein levels with hierarchical clustering. Data shown are representative of n=16 BP-IgG and n=9 control-IgG samples pooled from two independent experiments. **D)** Proteins decreased in BP patients with antibodies detectable only against BP230 vs. those with antibodies only detectable against BP180 normalized to control-IgG treated keratinocytes. Mann-Whitney U-test. **E)** BP-IgG induces histologic blistering in 3D HSE in the absence of immune cells. Mann-Whitney U-test. **F)** Direct immunofluorescence demonstrates deposition of anti-basement membrane IgG in BP-IgG but not control-IgG treated 3D HSE. NS – not significant. \**P* < 0.05, \*\**P* < 0.01. Scale bar = 100μm.

### BP-IgG Induces a Pro-inflammatory Transcriptional Response in Primary Human Keratinocytes

To validate these findings and to better understand the global effects of BP-IgG, we performed bulk RNA-seq on PHK treated with IgG from BP patients or controls. BP-IgG treated PHK demonstrated significant gene dysregulation relative to control-IgG treated PHK (**Figure 2A**, **Supplemental Data 2**) with notable dysregulation of numerous cytokines and chemokines (**Figure 2B**). Significant upregulation of *CXCL16, IL24*, and *TGFB1* corresponded with our protein findings, though *CXCL8* was notably downregulated at the RNA level in contrast to prior studies and protein levels^15^. We additionally identified significant dysregulation of complement components, S100 proteins, toll-like receptors, extracellular matrix (ECM) components, matrix metalloproteinases including *MMP9*, and skin barrier components (**Figure 2C**). Likewise, there was dysregulation of cytokeratins, including downregulation of *KRT10* and upregulation of *KRT17*, a marker of keratinocyte inflammation^42^. Several cluster of differentiation (CD) markers were significantly upregulated including *DPP4* (CD26), *CD14*, and *CD68*^43^ **(sFigure 1).** Gene ontology revealed enrichment in skin development, as well as chemotaxis, and T-cell modulation in response to BP-IgG (**sFigure 2**). To further validate our RNA-seq findings, we confirmed MMP9 upregulation by RT-PCR using a separate cohort of BP patients and PHK (**Figure 2D**).

**Figure 2:**
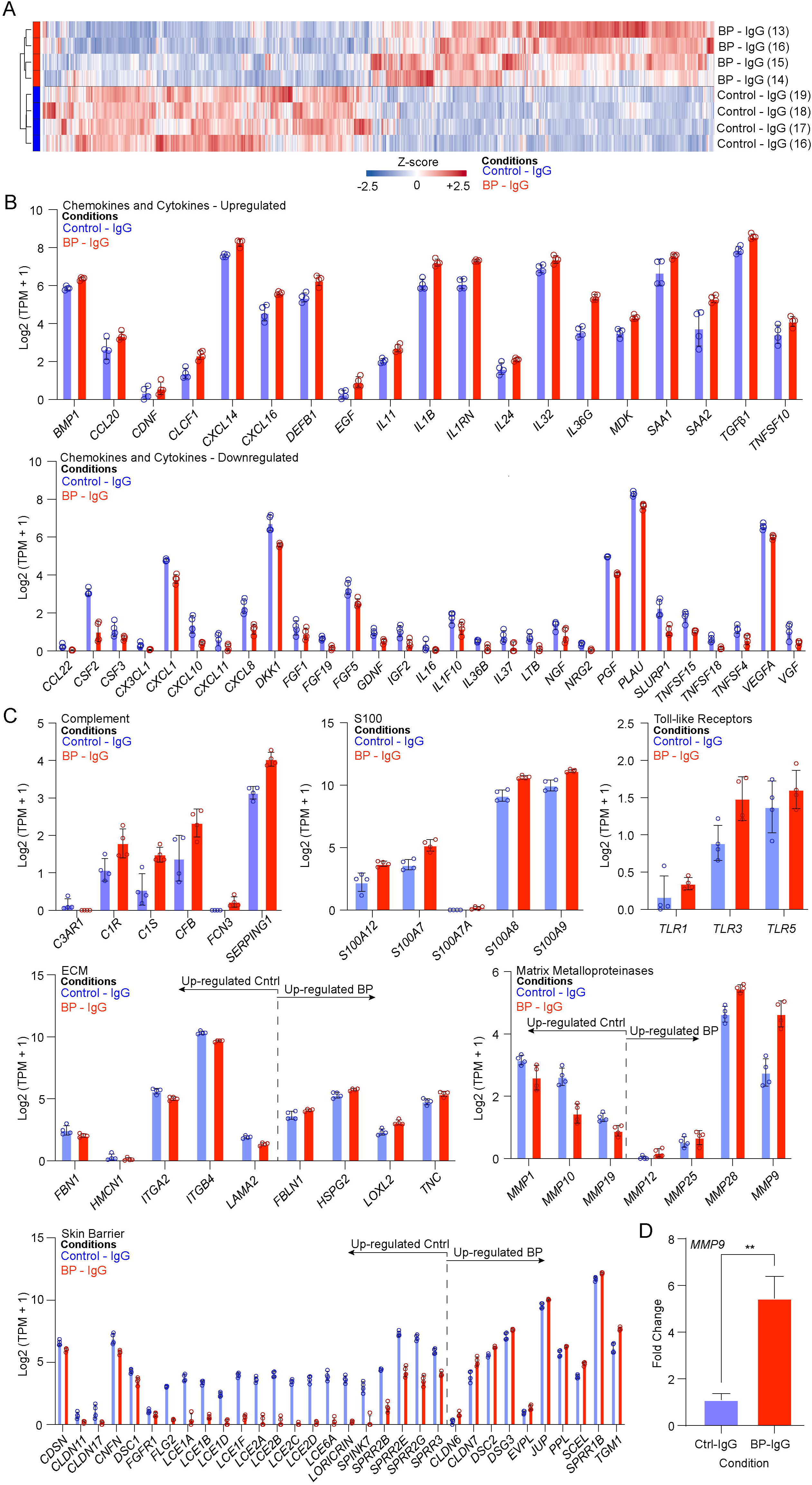
Whole transcriptome analysis of BP-IgG versus control-IgG-treated keratinocytes reveals dysregulation of numerous inflammatory and structural genes. **A)** Heat map of bulk RNA-seq from BP-IgG vs. control-IgG-treated primary human keratinocytes. **B)** Bar chart demonstrating normalized expression of significantly differentially expressed cytokines/chemokines, **C)** complement components, S100 family, toll-like receptors, extracellular matrix (ECM) components, matrix metalloproteinases, and skin barrier genes. Data shown is mean Log_2_(TPM+1) representative of n=4 per group. All data shown are significant at *P*_adj_ < 0.05. **D)** Validation of MMP9 upregulation by qPCR is shown as mean fold change ± SEM, representative of n=6 from a separate experiment. \*\**P* < 0.01.

### Blistered Epithelium in BP Mirrors BP-IgG Keratinocyte Culture Treatments

In light of these prior findings, we next aimed to determine how our BP-IgG-induced *in vitro* findings related to BP patient skin. First, we generated tissue microarrays from FFPE sections of patients with confirmed BP (**Figure 3A**). FFPE blocks were obtained from 20 patients with confirmed BP, as well as 20 samples of age-matched normal skin (NL), and spongiotic dermatitis with eosinophils (SD) as a control for eosinophil-rich inflammation. Patient demographics are summarized in supplementary table 2 (**sTable 2)**. Age did not significantly differ between BP and normal samples (i.e. 73.6 vs. 75.4, *P* = 0.52), while patients with spongiotic dermatitis were significantly younger than those with BP (i.e. 54.5 vs. 73.6, *P* < 0.01). Tissue microarrays were generated, sectioned, and placed on slides for spatial transcriptomics using GeoMx. To minimize the chance of including intraepithelial eosinophils or neutrophils, slides were stained for eosinophil peroxidase (EPX) or neutrophil elastase, respectively. Regions of interest (ROIs) were manually drawn onto the epithelium (**Figure 3B**). Blistering was scored as 0 (attached skin), 1 (fulcrum of a blister), or 2 (distant blister). Following quality control and normalization, spatial deconvolution was performed to confirm alterations in immune cell infiltration. No significant differences in cell abundance were observed, except for a decrease in melanocytes in the control groups (SD + NL) relative to BP (**sFigure 3)**.

**Figure 3:**
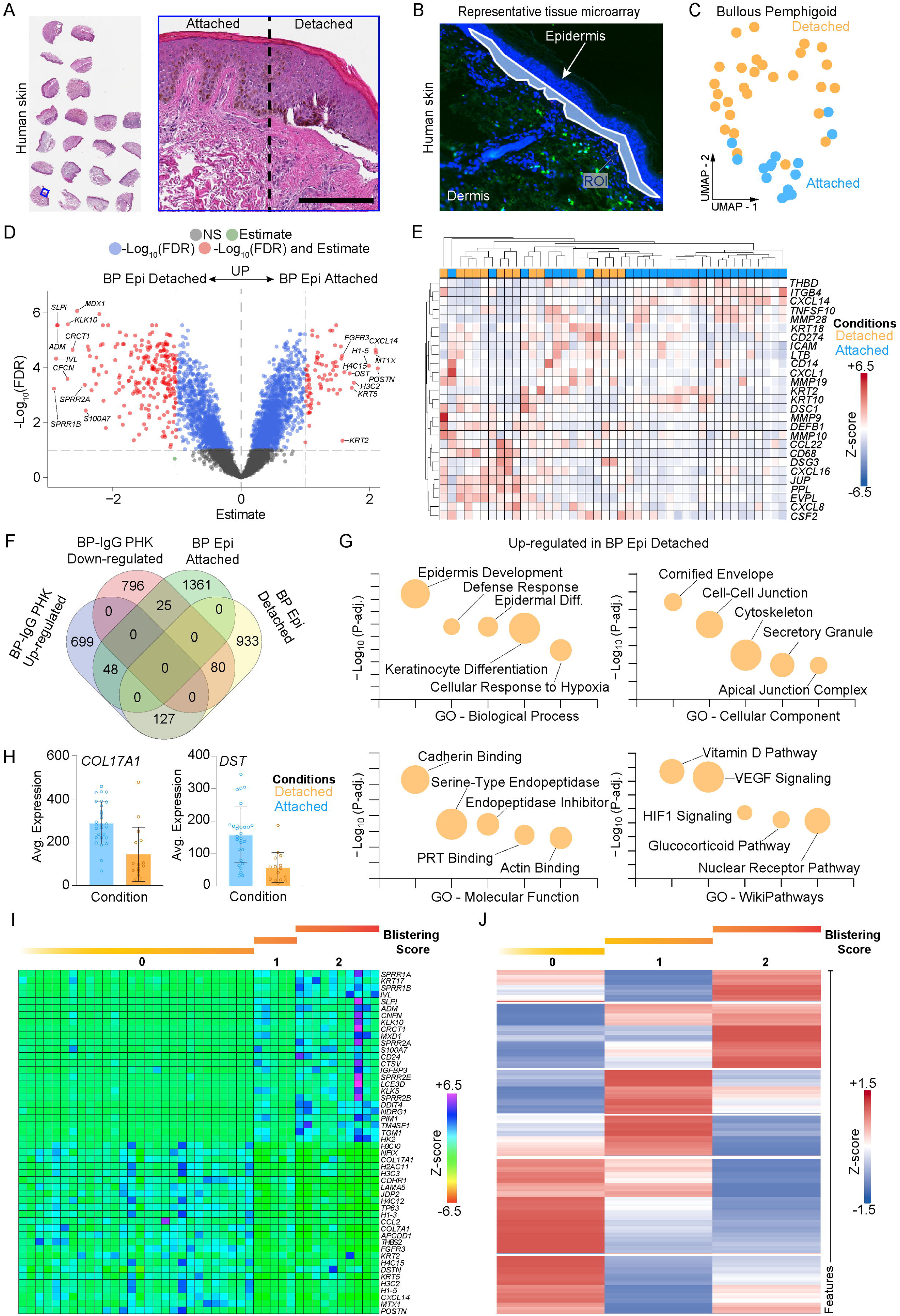
Spatial transcriptomics demonstrates a divergent transcriptome between blistered and attached skin in BP. **A)** Representative tissue microarray and distinction of attached vs. detached skin stained with H&E. **B)** Representative region of interest selection strategy (eosinophil peroxidase, EPX) shown in green; DAPI in blue. **C)** Two-dimensional UMAP plot demonstrating clustering of attached and detached epithelium in BP. **D)** Volcano plot of differentially expressed genes in detached vs. attached epithelia in BP. The top 10 differentially expressed genes sorted by *P*_adj_ are shown. **E)** Heat map comparing attached and detached gene expression from selected genes from BP-IgG-treated human keratinocyte experiments. **F)** Venn diagram of up-and down-regulated genes between BP-IgG vs. control-IgG treated primary human keratinocytes and detached vs. attached epithelium in BP. **G)** Dot plots of gene ontology (GO) and WikiPathways for detached vs. attached skin in BP. **H)** Differential gene expression of COL17a1 and DST in detached vs. attached BP skin. **I)** Gene expression trajectory of all BP patients in aggregate, and **J)** a single patient for whom all levels of blistering were available. Data shown is representative of 26 unique cores selected from n = 20 BP patients. Scale bar = 200μm. NS – not significant; ROI – region of interest.

We first questioned how detached (blistered) epithelia differed from attached epithelia in BP patient skin. Clustering and dimensionality reduction by UMAP revealed a clear distinction between the transcriptomes of attached and detached epithelia (**Figure 3C**). Differential gene expression analysis revealed 1,140 upregulated and 1,434 downregulated genes, defined as *P*_adj_ < 0.05 (**Figure 3D**, **Supplemental Data 3**). Numerous differentially expressed genes from BP-IgG treated keratinocytes were also noted in the detached BP epithelium, including small proline-rich proteins (SPRRs), kallikreins, *IVL, S100A7, CXCL16, CD68, MMP9*, and *JUP* (**Figure 3E**). While several genes were similarly upregulated or downregulated in spatial transcriptomics and primary cell experiments, others demonstrated opposite responses (**Figure 3F**, **Supplemental Data 4**). For example, *CXCL8* was significantly upregulated in patient skin, despite being downregulated in PHK experiments. Notably, spatial transcriptomics is less sensitive in identifying cytokine gene dysregulation, as described in previous studies on inflammatory skin disease, which may account for failure to identify differential expression of several cytokines and chemokines detected in 2D culture experiments such as IL-6 and IL-24 ^44^.

Gene ontology analyses further supported enrichment of keratinocyte differentiation pathways as well as VEGF and hypoxia-inducible factor-1 signaling (HIF-1a) pathways which presumably correspond with separation from the BMZ (**Figure 3G**). Upregulation of cornified envelope is also notable, as we have previously identified this as a response to dipeptidyl peptidase 4 (DPP4) inhibitors, a known trigger of BP^45^. Interestingly, *COL17A1* and *DST* were both significantly downregulated in detached skin, demonstrating a lack of a reparative mechanism and keratinocyte differentiation (**Figure 3H**).

We next sought to characterize the trajectory of gene expression as a function of blistering (**Figure 3I**). We aggregated patient transcriptomes from each level of blistering scored 0-2 to identify the top genes of each component, and plotted these as a heatmap, revealing a pseudo-trajectory from attached to blistered BP epithelium. This revealed increasing expression of differentiation markers such as SPRRs, kallikreins, and cathepsin L2, as well as late cornified envelope components. We then assessed this in a single patient in whom all ROIs of all 3 levels of blistering were present (**Figure 3J**), demonstrating a similar trajectory as a function of blistering.

### Attached BP Epithelium Demonstrates a Pro-inflammatory Transcriptome Shared with Spongiotic Dermatitis

Next, we questioned how BP epithelia differed from control skin to identify unique epithelial responses to BP-IgG. Based on our observations regarding the substantive differences between blistered and attached skin, we only considered attached skin to minimize potential bias from the blistering process for differentiated gene expression. Comparing BP epithelia to healthy control and spongiotic dermatitis as an inflammatory and eosinophil rich control in aggregate revealed minimal transcriptional alterations (**Figure 4A**), mostly driven by a lack of significant differences between BP epithelia and spongiotic dermatitis (**Figure 4B**). Clustering by UMAP demonstrated that spongiotic and BP attached epithelial mostly clustered together, with polarization between normal skin and detached BP epithelia (**Figure 4C**). Thus, in our comparative bioinformatic analyses, non-blistered epithelia in BP appears essentially indistinguishable from that of spongiotic dermatitis.

**Figure 4:**
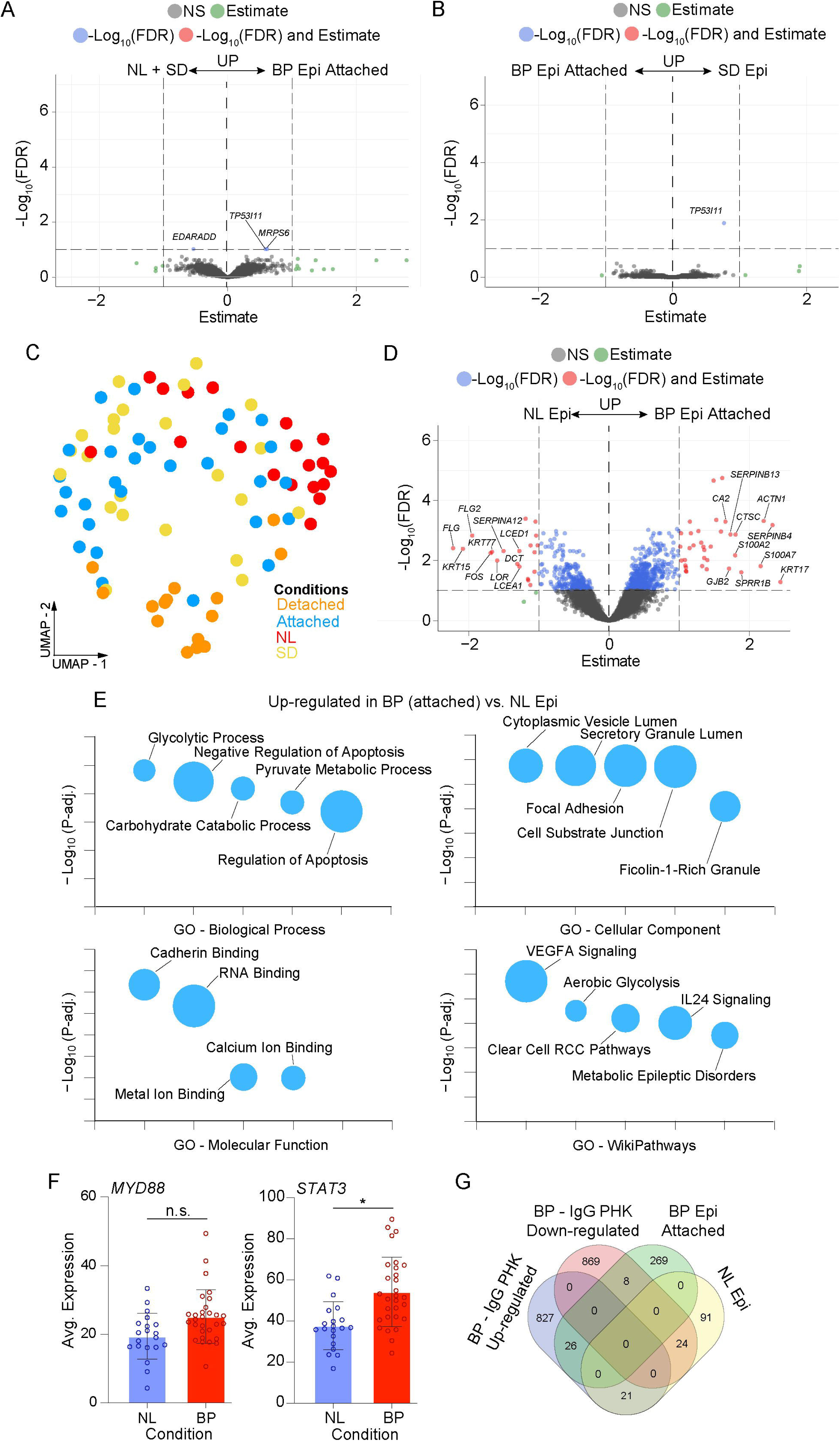
Spatial transcriptomics of attached BP skin and spongiotic dermatitis share numerous features distinct from normal skin. **A)** Volcano plot comparing BP epithelia with normal skin (NL) and spongiotic dermatitis (SD), revealing few differentially expressed genes along with **B)** BP vs. spongiotic dermatitis alone. **C)** Two-dimensional UMAP plot demonstrating significant overlap between BP (attached) and spongiotic dermatitis (SD) epithelial transcriptomes with polarization of BP (detached) and normal skin. **D)** Volcano plot demonstrating significantly differentially expressed genes between BP (attached) and normal (NL) skin defined as Log_2_F.C. > 0.5 and *P*_adj_ < 0.05. **E)** Gene ontology (GO) and WikiPathways bubble plots of BP (attached) vs. normal skin (NL). **F)** Upregulation of STAT3 (*P*_adj_ = 0.01) and MyD88 (*P*_adj_ = 0.12) expression in BP vs. normal skin. G) Venn diagram showing overlapping genes between BP (attached) vs. normal skin (NL) and BP-IgG vs. control-IgG-treated primary human keratinocytes. Data shown are representative of n=20 patients per cohort. NS – not significant.

When compared to normal epithelia only, attached BP epithelia demonstrated numerous differentially expressed genes (**Figure 4D**). Upregulation of several genes including *DSG3, KRT6A/KRT16*, as well as *S100A8/9, IL4R, VEGFA,* and *ADAM8* were noted, as well as downregulation of filaggrin genes (**Supplemental Data 5**). Significantly enriched pathways in BP skin relative to normal skin include cytoplasmic vesicle lumen, consistent with expected macropinocytosis induced by anti-BP180-IgG *in vitro*, as well as VEGFA signaling (**Figure 4E**). Interestingly, we noted significant upregulation of STAT3 in attached BP epithelia versus normal skin, with a trend towards increased MyD88 (**Figure 4F**). We then investigated the overlap between upregulated genes in BP epithelia relative to normal, and those differentially expressed in PHK experiments (**Figure 4G**, **Supplemental Data 6**). Notably, VEGFA and ADAM8 were downregulated in BP-IgG-treated keratinocytes yet upregulated in BP epithelium, suggesting indirect mechanisms of upregulation.

### Comparison of Keratinocyte Transcriptomes by pseudo-bulk scRNA-seq

We next utilized the recently published database by Liu et al, which included single cell transcriptomes from the lesion skin of 8 BP patients and 5 healthy controls^46^. In total we assessed 27,591 keratinocyte transcriptomes (**Figure 5A, B**). Keratinocytes were divided as basal, suprabasal, and granular based on expression of canonical markers KRT14, KRT10, and IVL respectively, with a comparable proportion between samples (**sFigure 4**). We next performed pseudo-bulk differential gene expression of keratinocytes in aggregate (**Figure 5C**, **Supplemental Data7**), which similarly reflected differential gene expression of basal keratinocytes between BP and controls (**Supplemental Data 8)**. These findings mirrored spatial transcriptomics experiments, with upregulation of several inflammatory and differentiation markers including ADAM8, C1s, DPP4, KRT6A/KRT16, IL4R, MyD88, TGFBI, and VEGFA (**Figure 5D**).

**Figure 5:**
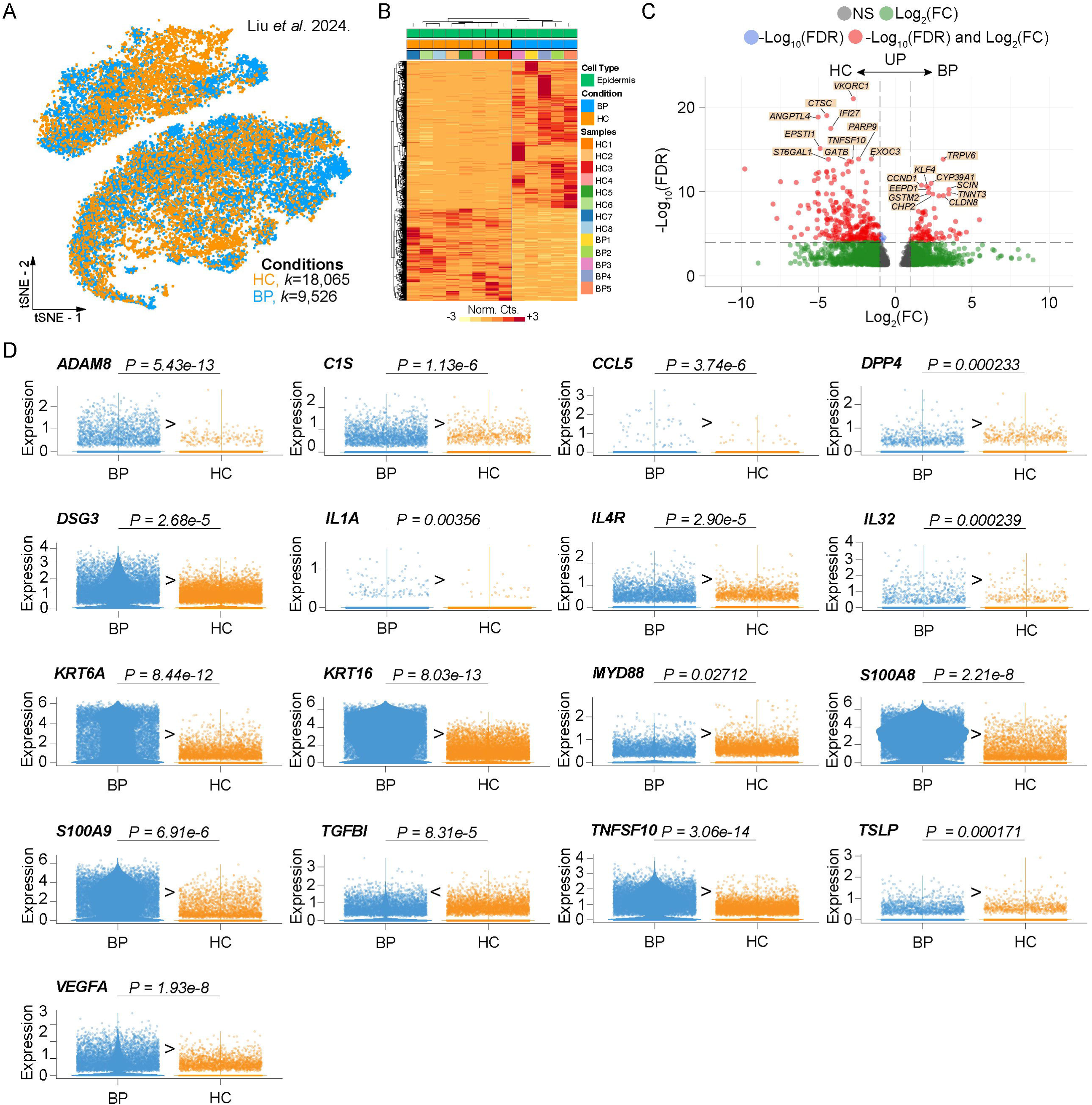
scRNA-seq of BP versus normal keratinocytes: **A)** t-SNE feature plot demonstrating bioinformatically gated keratinocyte transcriptomes from BP vs. healthy control (HC) patient skin. **B)** Heat map of BP vs. HC skin. **C)** Volcano plot with the top 10 differentially expressed genes between all keratinocytes in BP patients vs. HC skin cohort. **D)** Violin plots of key differentially expressed between BP keratinocytes and HC keratinocytes using pseudobulk analysis of scRNA-seq reveals significant dysregulation of several inflammatory, differentiation, and protease markers. NS – not significant.

### Keratinocyte MyD88 Regulates Much of the Inflammatory Response to BP-IgG

Given the upregulation of numerous MyD88 and NF-κB regulated cytokines in PHK experiments^39, 47–50^, the upregulation of STAT3^51^ in BP epithelia, and upregulation of MyD88 in scRNA-seq data, we sought to further investigate MyD88 regulation of the keratinocyte. We utilized the MyD88 Crispr/Cas9 mediated knockout (MyD88 KO) nTERT cells along with transfection control nTERT^39^ cells, repeating treatment with BP-IgG or control-IgG.

First, we performed bulk-RNA-seq of BP-IgG treated nTERT and MyD88 KO cells, as well as control-IgG treated MyD88 KO cells (**Figure 6A**, **Supplemental Data 9**). Principal component analysis (PCA) demonstrated distinct clustering of each respective cohort (**Figure 6B**). Bulk RNA-seq revealed reduced upregulation of numerous inflammatory markers in response to BP-IgG in MyD88 KO relative to BP-IgG treated control cells, including *IL1B, IL24, IL36G, TSLP, CXCL8, MMP1, CSF3,* and *S100A7/8/9/P* (**Figure 6D**). Barrier alarmin molecules were additionally down regulated in MyD88 KO in response to BP-IgG including *CLDN1/7/14/23, KRT6b*, and *KRT16* (**sFigure 5**). Bulk RNA-seq of BP-IgG vs control-IgG in MyD88 KO still demonstrated several findings from PHK (**Supplemental Data 10**), including upregulation of *MMP9* and *IL24*.

**Figure 6:**
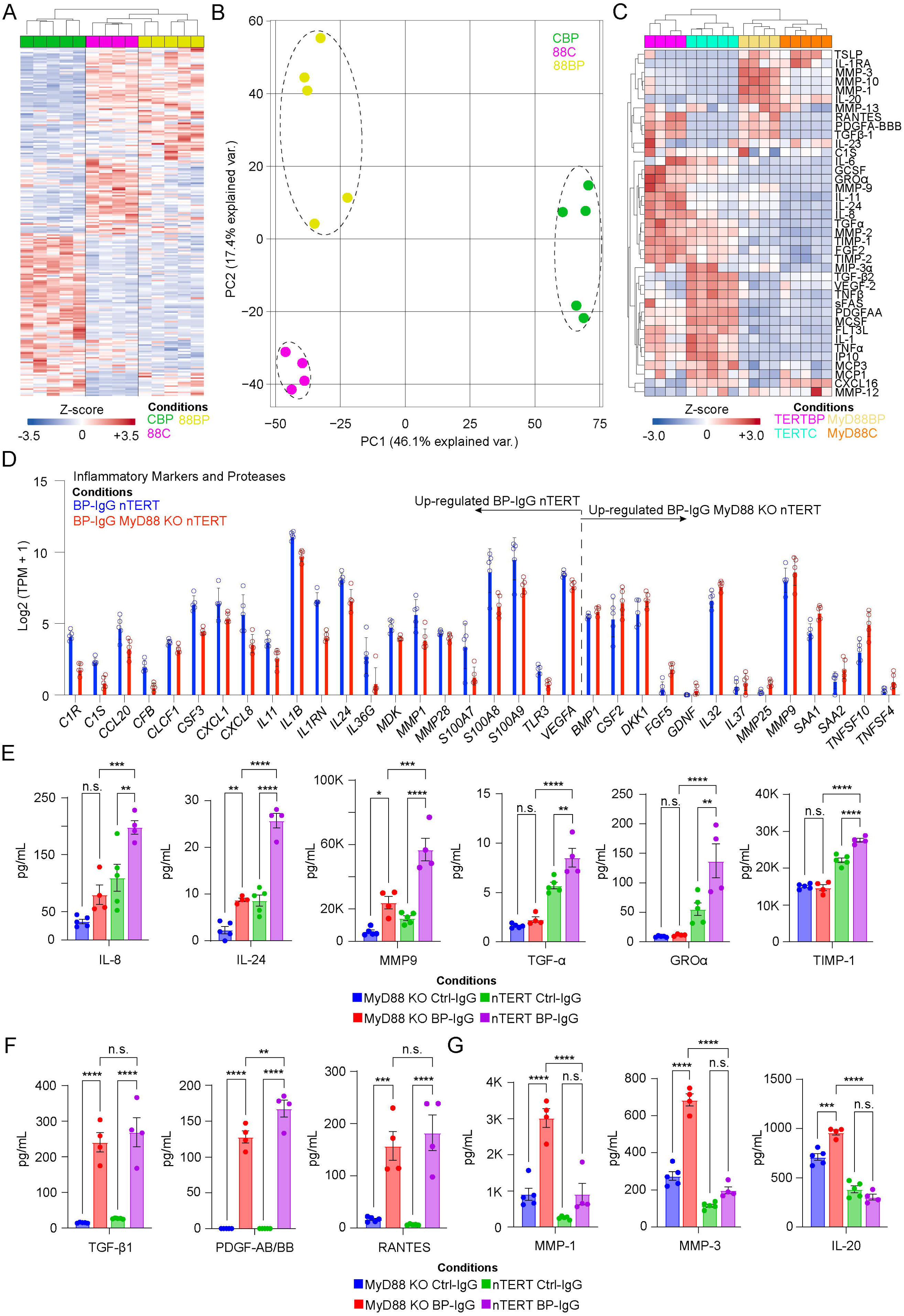
MyD88 regulates a number of responses to BP-IgG in keratinocytes. **A)** Heat map of gene expression from BP-IgG-treated control (CBP) or MyD88-deficient nTERT cells (88BP) and control-IgG-treated MyD88-deficient nTERT cells (88C). **B)** Two-dimensional PCA plot demonstrates clustering of samples, supporting significant transcriptional changes due to MyD88 deficiency. **C)** Heat map of supernatant proteins from BP-IgG-vs. control-IgG-treated MyD88 KO nTERT (MyD88BP, MyD88C) and control nTERT cells (TERTBP TERTC). **D)** Bar chart demonstrating selected genes induced by BP-IgG that are affected by MyD88 deficiency. All data shown are significant at *P*_adj_ < 0.05. **E)** MyD88 knockout blunts protein expression of IL-8, IL-24, MMP-9, TGFα, GROα, and TIMP-1. **F)** Increased protein expression of TGF-β1, PDGF-AB/BB, and RANTES occurs regardless of MyD88 deficiency. **G)** MyD88 KO results in a relative increase in MMP-1, MMP-3, and IL-20 responses. Data shown are representative of n=4-5 per cohort shown as mean ± SEM. ANOVA with Tukey’s test. NS – not significant. \**P* < 0.05, \*\**P* < 0.01, \*\*\**P* < 0.001, and \*\*\**P* < 0.0001.

Analysis of supernatant cytokines and chemokines demonstrated similar clustering based on supernatant protein expression (**Figure 6C**, **Supplemental Data 11**). As suspected, IL-8, IL-24, MMP9, TGFα, GROα, and TIMP-1 were significantly blunted in MyD88 KOs (**Figure 6E**). In contrast, TGF-β1, PDGF-AB/BB, and RANTES were upregulated by BP-IgG treatment independent of MyD88 status (**Figure 6F**), while BP-IgG induced a more robust MMP-1, MMP-3, and IL-20 response in MyD88 KO than control cells (**Figure 6G**). Notably G-CSF was undetectable, while IL-1α, IP-10, GROα, M-CSF and TNFα were significantly reduced in MyD88 KO cells regardless of IgG treatments (**sFigure 6**).

### Keratinocyte Dependent MyD88 Knockout Decreases Disease Severity in an Experimental Model of Bullous Pemphigoid

In light of these findings, we questioned whether keratinocyte-specific MyD88 knockout could limit pathology in an *in vivo* model induced by anti-COL17 IgG. To test this, we next generated *Krt14*-dependent MyD88 knockouts (*Krt14*-Cre^+^;*Myd88*^fl/fl^) as previously described^52^, using *Krt14*-Cre^-^;*Myd88*^fl/fl^ mice as controls. Rabbit anti-mouse COL17^NC14–1^ was injected into age-and sex-matched mice over a period 2 weeks as previously described^36^. *Krt14*-Cre^+^;*Myd88^fl/fl^* mice demonstrated a significantly decreased affected body surface area relative to controls (**Figure 7A, B**).

**Figure 7:**
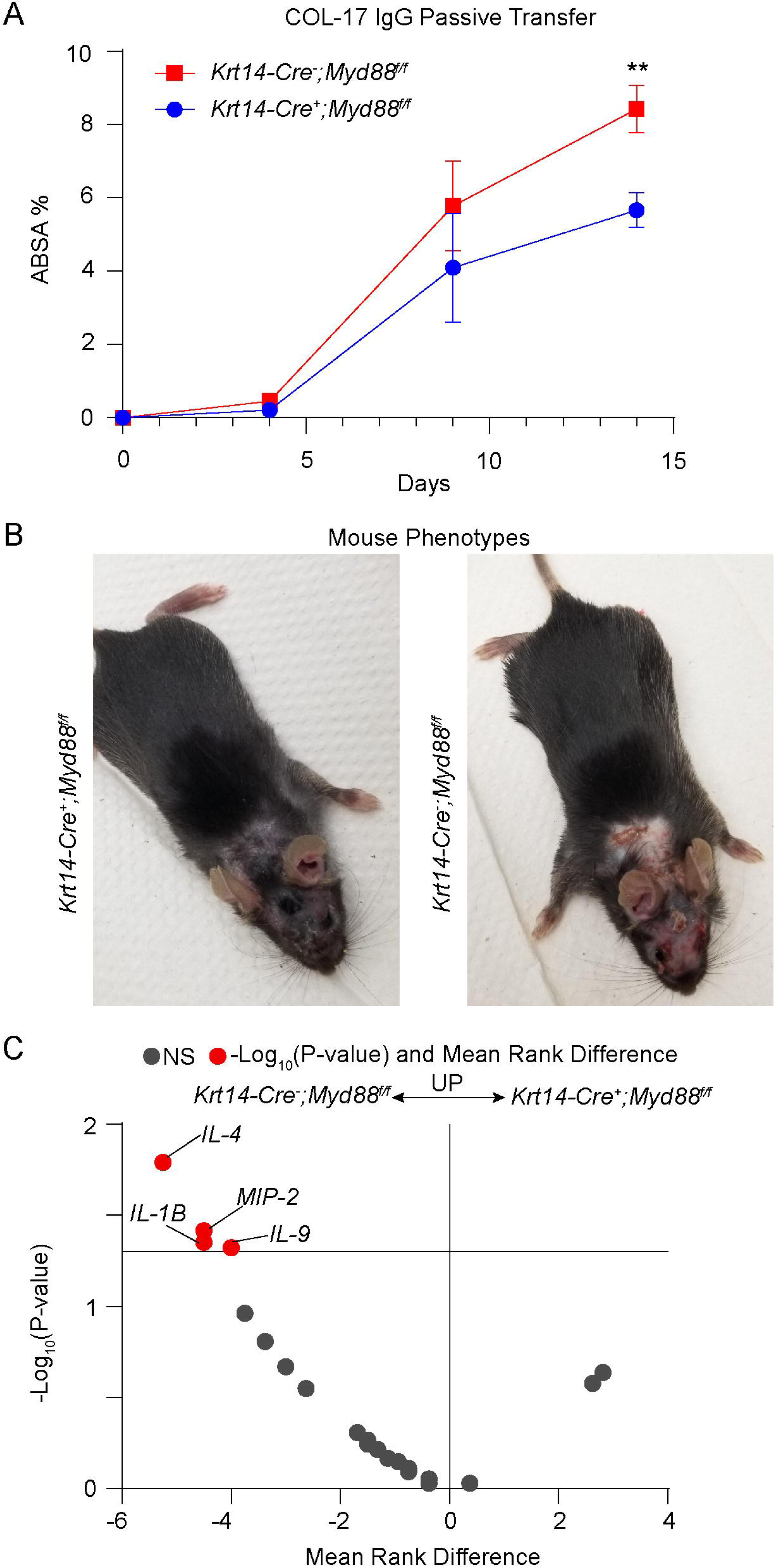
*Krt14*-dependent *Myd88* deficiency decreases disease severity in BP, reducing inflammatory cytokines. **A)** Decreased affected body surface area score (ABSA) is seen at day 14 in *Krt14-Cre^+^;Myd88^fl/fl^*. **B)** Representative phenotype of *Krt14-Cre^+^;Myd88^fl/fl^* and *Krt14Cre^-^-Myd88^fl/fl^*mice. **C)** Volcano plot demonstrating decrease serum levels of IL-1β, IL-4, MIP-2, and IL-9 in keratinocyte-dependent *Myd88* knockouts relative to controls. Data shown are mean ± SEM from n=6-8 mice pooled from two independent experiments. Mann Whitney U-test. NS – not significant. \*\**P* < 0.01.

### Keratinocyte Dependent MyD88 Deficiency Decreases Key Circulating Type 2 Cytokines and Chemokines

As we demonstrated an overall decrease in disease severity as a result of MyD88 knockout in murine keratinocytes, we hypothesized that this decreased keratinocyte response would translate to a decreased systemic inflammatory response. As such, we performed a 32-plex cytokine/chemokine multiplex immunoassay of serum from *Krt14*-Cre^+^*Myd88^fl/fl^* or controls. Notably, serum from keratinocyte-dependent MyD88 knockout mice demonstrated a significant decrease in serum IL-1β, IL-4, IL-9, and MIP-2 levels relative to controls (**Figure 7C**).

## Discussion

Our data reveal a novel role of keratinocytes as an orchestrator of the inflammatory and proteolytic response seen in BP. Both transcriptomic and large-scale protein assays demonstrated a proinflammatory response in primary keratinocytes treated with BP-IgG that was similar to gene expression in blistered epithelium of BP patients, providing evidence that these autoantibody-dependent effects may be contribute to disease pathology. Non-blistered epithelium in BP, by contrast, is not significantly transcriptionally different from the epithelium in spongiotic dermatitis. This builds on prior evidence that bullous and non-bullous phenotypes demonstrate different inflammatory responses^53, 54^ Many of the genes upregulated in BP-IgG-treated primary keratinocytes are known to be regulated by MyD88 and/or NFκB, and we found that knockout of keratinocyte MyD88 *in vitro* and *in vivo* results in blunting of many of these factors, decreasing clinical disease severity, as well as serum cytokines including IL-1β, IL-4, and IL-9.

Whether keratinocytes are a bystander in BP, or the driver of systemic response is not clear^5, 6^. We posit that keratinocytes are active drivers of disease in response to BP-IgG. Prior studies demonstrating keratinocyte upregulation of IL-6 and IL-8 in response to anti-BP180 antibodies suggest more than a bystander role in the inflammatory response. The occurrence of this with both IgG and IgE autoantibodies additionally highlights independence from the immune cell Fc receptor^13^. As keratinocytes release numerous pro-inflammatory markers, proteases, and complement components, this provides further support of a keratinocyte-derived systemic response. The translation of these 2D culture findings to *in vivo* pathogenicity remains unclear. The efficacy of whole-body topical corticosteroid treatment relative to oral corticosteroids does provide some insight into skin as an inflammatory organ^55^. While this may be a result of direct action on skin-infiltrating leukocytes, topical corticosteroids may also directly inhibit inflammatory responses of keratinocytes. The upregulation of C1 complex components’ seen in BP-IgG treated keratinocytes also raises the question as to whether the keratinocyte pathologically contributes towards a complement-dependent response^56^. Notably, a C1s antibody was shown to completely block complement pathway activation using serum from BP patients *in vitro*^57^. Thus, the observation of release of C1s from keratinocytes induced by BP-IgG in serum free 2D conditions, could play a major role in the complement-dependent response in BP.

The blunting of systemic IL-4 and IL-9 increases in *Krt14*-specific *Myd88* KO mice may be particularly clinically relevant. The anti-IL4Rα antibody dupilumab is currently under investigation for the treatment of BP, with numerous retrospective studies demonstrating its efficacy^58–61^. Likewise, IL-9 has recently been shown to correlate with clinical responses in BP^62^. Th9 cells have been identified in skin samples of BP and are able to drive eosinophil-rich and type 2 inflammatory responses in numerous other diseases^62–68^. The expression of IL-1β and TGF-β1 seen in BP-IgG-treated keratinocytes is also noteworthy, as when combined with IL-4, these can drive the differentiation of naïve T cells to Th9 cells^63, 69^. As neither IL-4 nor IL-9 is expressed by keratinocytes, our finding of decreased serum levels in *Krt14*-dependent *Myd88* knockout mice highlights the interaction between keratinocyte-mediated inflammation as a response to BP-IgG, and development of Th2/Th9 responses thereafter. The role of other T-cell stimulatory factors identified in BP-IgG-treated keratinocytes such as CTACK^70^, IL-24^71^, IL-6^72^, CXCR16^73^, and RANTES^74, 75^ warrants further investigation.

VEGFA was also highly upregulated in BP epithelium in our spatial transcriptomics analysis, along with NRP1 which is required for VEGFA endothelial function^76^. While numerous factors can induce VEGFA, previous work has demonstrated that keratinocytes treated with IL-9 upregulated VEGFA. Circulating VEGFA has also been associated with an increased risk of venous thromboembolism^77, 78^, a known complication of BP thought to occur due to local coagulation factors^79–81^. This finding aligns with previous studies identifying elevated VEGFA protein in BP blister fluid^82^, serum^83^, and skin^84^. As VEGF-A was not directly induced by BP-IgG on keratinocytes, this would indicate an intermediary stimulus to induce upregulation of certain factors, such as IL-9.

Keratinocyte-derived proteases should also be considered in the context of granulocyte chemotaxis. While MMP-9 is capable of cleaving BP180, it also appears to regulate eosinophil chemotaxis to the skin in BP. A recent study utilizing an IgE model of BP demonstrated decreased eosinophil infiltration in *Mmp9* knockout mice^85^. We have also identified ADAM8 in BP epithelium which can also act as an eosinophil chemoattractant in other disease models^86^. Notably, we did not detect ADAM8 in BP-IgG treated PHK supernatants, again likely indicating indirect stimulation on human skin. ADAM8 has been shown to be essential in the development of experimental asthma^87^ and its inhibition has been shown to decrease eosinophil and Th2 lymphocyte infiltration in the lungs^88^. It is thought to function through cleavage of CD23 into a soluble form^89^. Interestingly, ADAM8 was found to correlate with airway allergen induced pneumonitis, but not with blood (drug-induced) pneumonitis, thus further implicating epithelial-immune interactions^89^. There is only limited data pertaining to ADAM8 in the skin, with a study demonstrating transgenic animals exhibit more severe oxazolone-induced cutaneous hypersensitivity^90^. Further analysis in the context of BP is needed, especially since high levels of sCD23 were found in BP blister fluid compared to suction blisters^91^.

Despite the histological abundance of eosinophils in BP skin, there was a notable lack of major eosinophil chemotactic factors induced by BP-IgG, aside from MMP-9 and RANTES^92^. While eotaxin-1 is expressed in BP skin as well as in spongiotic dermatitis^35, 93^, basal expression on keratinocytes is decreased when treated with BP-IgG. While RANTES also acts as a chemotactic factor for eosinophils, its expression on BP has also been questioned, as it was not detected in an immunohistochemistry study of BP skin versus normal skin^94^ or in BP serum^95^. Thus, we hypothesize that keratinocytes release factors that indirectly drive eosinophil chemotaxis. However, this warrants further investigation.

We additionally identified upregulation of DPP4 in both BP-IgG-treated keratinocytes and BP skin. As DPP4 inhibitors lead to a significantly increased risk of developing BP^96^, this finding appears counterintuitive. DPP4 is an exopeptidase that can act on numerous secreted factors. DPP4 can drive Th1/Th17 proliferation^96^, while increased number of Th17 cells have been described in early BP skin lesions^97^. Soluble DPP4 levels have been noted to negatively correlate with systemic inflammation which contrasts with our findings of upregulated DPP4 in the setting of release of inflammatory molecules^98^. DPP4 can, however, also cleave RANTES which may account for prior findings of a lack of detectable protein on human skin^99^, as well as cleave eotaxin^100^. Inhibition of DPP4 on keratinocytes leads to upregulation of the late cornified envelope and cytoskeletal remodeling, but an absence of inflammatory or proteolytic findings^45, 101^.Thus, whether oral DPP4 inhibitors act on keratinocyte-derived DPP4 induced by BP-IgG, or through inhibition of upstream T-cell responses, remains unanswered.

BP antibodies appear to also exert a significant effect on keratinocyte differentiation. While bulk RNA-seq identified a decrease in ITGB4 expression, we identified significant downregulation of numerous hemidesmosomal genes on detached BP epithelium including *COL17A1, DST, ITGB4, ITGA3*, and the recently characterized *LAMB4* which serves as the autoantigen in p200 pemphigoid^102, 103^. Decreased *KRT5* and *KRT14* along with increases in granular layer markers such *as IVL, TGM1, FLG,* and *LCE3D* further point towards keratinocyte differentiation. We have previously demonstrated that blockade of the laminin-332 and integrin α6β4 interaction was sufficient to induce blistering and result in keratinocyte differentiation via protein kinase C and NOTCH^21^. Thus, BP appears to demonstrate a similar phenomenon, whereby autoantibody-induced blistering is perpetuated by a loss of differentiation with loss of hemidesmosomal gene expression.

Our study has several limitations. Total IgG rather than BP180 or BP230 specific IgG was extracted from serum. Thus, the impact of alternative autoantigens or differences in the dose of pathogenic autoantibodies cannot be ruled out. We did not, however, identify differences amongst most markers assessed in patients with BP180 versus BP230 antibodies. We also lacked a sufficient sample size of patients receiving DPP4 inhibitors, which are described to target different epitopes of BP180 and have a less inflammatory response^104^, to perform a subset analysis. Likewise, due to differences in diagnostic tests and batch effects between experiments, we could not pool samples to perform an adequate correlation analysis of indirect immunofluorescence titers or BP180/BP230 levels with protein or gene expression. Spatial transcriptomics provides a pseudo-bulk-transcriptomic view that lacks single-cell resolution and prevents the identification of single epithelial and immune cells and their respective interactions. This was partially supplemented by employing and analyzing a scRNA-seq dataset of BP patients and controls, though these were from different patients. This dataset likewise revealed skin lesions with different cellular composition from prior studies identifying lymphocyte and eosinophil-predominant inflammation^105^, though this was presumably due to technical challenges of scRNA-seq in capturing granulocyte transcriptomes^106^. Nonetheless, by clustering blistered versus attached BP skin, we were able to gain significant insight into transcriptional differences and perform a pseudo-trajectory analysis. This demonstrated a critical role of keratinocyte differentiation as a function of skin separation. As 19 of 20 patients demonstrated C3 deposition on direct immunofluorescence, we were likewise unable to perform a subanalysis based on the presence of complement. As blood and skin samples were also not matched, we could not compare individual paired keratinocyte responses and skin expression of key inflammatory markers. Lastly, given we noted significantly decreased IL-1α, G-CSF, IP-10, M-CSF and TNF-α in MyD88-deficient keratinocytes we cannot rule out that these were the critical factors resulting in decreased disease severity *in vivo* and type 2 response rather than blunted IL-24 or MMP9 expression. Likewise, as CXCL8 is not expressed in mice, it is unclear to what extent cytokines and chemokines are blocked in mouse MyD88-deficient keratinocytes relative to human keratinocytes. Still, given a decrease in disease severity *in vivo*, these observations nonetheless implicate keratinocytes in coordinating the systemic response to BP-IgG.

In conclusion, we demonstrate that the keratinocyte plays an active role in the response to autoantibodies in BP. IgG autoantibodies induce an inflammatory response with the release of proteases. These antibodies are sufficient to induce blistering independent of granulocytes, leading to keratinocyte differentiation and loss of expression of hemidesmosomal genes. This keratinocyte response is regulated in part through MyD88 signaling, resulting in decreased disease severity and a reduction of several cytokines and proteases. Thus, our study points to the important contributory role of keratinocytes to the pathogenesis of BP.

## Abbreviations

BMZ: basement membrane zone
BP: bullous pemphigoid
COL17A1: collagen type XVII alpha 1 chain
COL17: collagen type XVII
CRISPR: Clustered Regularly Interspaced Short Palindromic Repeats
DPP4: dipeptidyl peptidase 4
ELISA: enzyme-linked immunosorbent assay
FcR: Fc receptor
FDR: false discovery rate
FFPE: formalin-fixed paraffin,-embedded
HSE: human skin equivalent
Krt14: Keratin 14
PCA: Principal component analysis
PHK: Primary Adult Human Keratinocytes
ROI: region of interest
RT-PCR: Reverse Transcription Polymerase Chain Reaction
RIN: RNA integrity number
ROI: Region of interest
scRNA-seq: single-cell RNA sequencing
SPRRs: Small Proline-Rich Protein
TMA: Tissue Microarray
TSLP: thymic stromal lymphopoietin
UMAP: Uniform Manifold Approximation and Projection

## Acknowledgements

We thank UIC Research Histology and Research Tissue Imaging cores for help with tissue samples sectioning, staining, and imaging of the slides.

**sFigure 1:** Bar chart of significantly dysregulated cytokeratins and cluster of differentiation markers in BP-IgG vs. control-IgG-treated keratinocytes. Data shown are mean Log_2_(TPM+1), representative of n=4 per group. All data shown have *P*_adj_ < 0.05.

**sFigure 2:** Bubble plot of gene ontology and WikiPathways enriched in BP-IgG vs. control-IgG-treated keratinocytes. The top 5 **A)** up-regulated and **B)** down-regulated pathways using gene ontology for biologic process, molecular function, cellular component, and WikiPathways are shown.

**sFigure 3:** Spatial deconvolution from BP, normal (NL), and spongiotic dermatitis (SD) gene sets. Data demonstrate a significant alteration in melanocyte abundance in BP vs. NL and SD, but no significant alteration in inflammatory cell estimation after false discovery rate correction. NS – not significant.

**sFigure 4: A)** t-SNE plot of keratinocyte from BP and HC. **B)** Relative cell abundance of *KRT14*, *KRT10*, or *IVL* expressing keratinocytes between BP skin and HC. **C)** Feature plot demonstrating *KRT14*, *KRT10*, and *IVL* expression.

**sFigure 5:** Bar charts of differentially expressed genes in skin barrier regulation in MyD88 deficient or control nTERT cells treated with IgG from patients with BP. Data shown are Log_2_(TPM+1) from n=4 patients per group. All data shown have *P*_adj_ < 0.05.

**sFigure 6:** Bar chart demonstrating significant downregulation of IL-1α, IP-10, GROα, M-CSF, and TNF-α in MyD88 deficient cells irrespective of IgG treatment(s). Data shown are representative of n=4-5 per cohort, shown as mean ± SEM. ANOVA with Tukey’s test. \**P* < 0.05, \*\**P* < 0.01, \*\*\**P* < 0.001, **** *P* < 0.0001. *n.s. – not significant*.

## References

1. Amber KT, Murrell DF, Schmidt E, Joly P, Borradori L. Autoimmune Subepidermal Bullous Diseases of the Skin and Mucosae: Clinical Features, Diagnosis, and Management. Clin Rev Allergy Immunol 2018; 54:26–51.

2. Cole C, Borradori L, Amber KT. Deciphering the Contribution of BP230 Autoantibodies in Bullous Pemphigoid. Antibodies (Basel) 2022; 11.

3. Hiroyasu S, Ozawa T, Kobayashi H, Ishii M, Aoyama Y, Kitajima Y, et al. Bullous pemphigoid IgG induces BP180 internalization via a macropinocytic pathway. Am J Pathol 2013; 182:828–40.

4. Bao L, Perez White BE, Chang RC, Li J, Vanderheyden K, Verheesen P, et al. FcRn inhibition reduces bullous pemphigoid anti-basement membrane zone IgG deposition and blistering in 3D human skin equivalents. J Invest Dermatol 2024.

5. Cole C, Vinay K, Borradori L, Amber KT. Insights Into the Pathogenesis of Bullous Pemphigoid: The Role of Complement-Independent Mechanisms. Front Immunol 2022; 13:912876.

6. Papara C, Karsten CM, Ujiie H, Schmidt E, Schmidt-Jiménez LF, Baican A, et al. The relevance of complement in pemphigoid diseases: A critical appraisal. Front Immunol 2022; 13:973702.

7. Schulze FS, Beckmann T, Nimmerjahn F, Ishiko A, Collin M, Köhl J, et al. Fcγ receptors III and IV mediate tissue destruction in a novel adult mouse model of bullous pemphigoid. Am J Pathol 2014; 184:2185–96.

8. Amber KT, Valdebran M, Kridin K, Grando SA. The Role of Eosinophils in Bullous Pemphigoid: A Developing Model of Eosinophil Pathogenicity in Mucocutaneous Disease. Front Med (Lausanne) 2018; 5:201.

9. Jones VA, Patel PM, Amber KT. Eosinophils in bullous pemphigoid. Panminerva Med 2021; 63:368–78.

10. Chiorean RM, Baican A, Mustafa MB, Lischka A, Leucuta DC, Feldrihan V, et al. Complement-Activating Capacity of Autoantibodies Correlates With Disease Activity in Bullous Pemphigoid Patients. Front Immunol 2018; 9:2687.

11. Karsten CM, Beckmann T, Holtsche MM, Tillmann J, Tofern S, Schulze FS, et al. Tissue Destruction in Bullous Pemphigoid Can Be Complement Independent and May Be Mitigated by C5aR2. Front Immunol 2018; 9:488.

12. Freire PC, Muñoz CH, Stingl G. IgE autoreactivity in bullous pemphigoid: eosinophils and mast cells as major targets of pathogenic immune reactants. Br J Dermatol 2017; 177:1644–53.

13. Messingham KN, Crowe TP, Fairley JA. The Intersection of IgE Autoantibodies and Eosinophilia in the Pathogenesis of Bullous Pemphigoid. Front Immunol 2019; 10:2331.

14. van Beek N, Lüttmann N, Huebner F, Recke A, Karl I, Schulze FS, et al. Correlation of Serum Levels of IgE Autoantibodies Against BP180 With Bullous Pemphigoid Disease Activity. JAMA Dermatol 2017; 153:30–8.

15. Schmidt E, Reimer S, Kruse N, Bröcker EB, Zillikens D. The IL-8 release from cultured human keratinocytes, mediated by antibodies to bullous pemphigoid autoantigen 180, is inhibited by dapsone. Clin Exp Immunol 2001; 124:157–62.

16. Schmidt E, Reimer S, Kruse N, Jainta S, Bröcker EB, Marinkovich MP, et al. Autoantibodies to BP180 associated with bullous pemphigoid release interleukin-6 and interleukin-8 from cultured human keratinocytes. J Invest Dermatol 2000; 115:842–8.

17. Messingham KN, Srikantha R, DeGueme AM, Fairley JA. FcR-independent effects of IgE and IgG autoantibodies in bullous pemphigoid. J Immunol 2011; 187:553–60.

18. Ruan Y, Xu C, Zhang T, Zhu L, Wang H, Wang J, et al. Single-Cell Profiling Unveils the Inflammatory Heterogeneity within Cutaneous Lesions of Bullous Pemphigoid. J Invest Dermatol 2024.

19. Hu Z, Zheng M, Guo Z, Zhou W, Zhou W, Yao N, et al. Single-cell sequencing reveals distinct immune cell features in cutaneous lesions of pemphigus vulgaris and bullous pemphigoid. Clin Immunol 2024; 263:110219.

20. Bao L, Li J, Solimani F, Didona D, Patel PM, Li X, et al. Subunit-Specific Reactivity of Autoantibodies Against Laminin-332 Reveals Direct Inflammatory Mechanisms on Keratinocytes. Front Immunol 2021; 12:775412.

21. Bao L, Perez White BE, Li J, Patel PM, Amber KT. Gene expression profiling of laminin α3-blocked keratinocytes reveals an immune-independent mechanism of blistering. Exp Dermatol 2022; 31:615–21.

22. Jones VA, Patel PM, Gibson FT, Cordova A, Amber KT. The Role of Collagen XVII in Cancer: Squamous Cell Carcinoma and Beyond. Front Oncol 2020; 10:352.

23. Tuusa J, Kokkonen N, Tasanen K. BP180/Collagen XVII: A Molecular View. Int J Mol Sci 2021; 22.

24. Zhang Y, Hwang BJ, Liu Z, Li N, Lough K, Williams SE, et al. BP180 dysfunction triggers spontaneous skin inflammation in mice. Proc Natl Acad Sci U S A 2018; 115:6434–9.

25. Hurskainen T, Kokkonen N, Sormunen R, Jackow J, Löffek S, Soininen R, et al. Deletion of the major bullous pemphigoid epitope region of collagen XVII induces blistering, autoimmunization, and itching in mice. J Invest Dermatol 2015; 135:1303–10.

26. Saraiya A, Yang CS, Kim J, Bercovitch L, Robinson-Bostom L, Telang G. Dermal eosinophilic infiltrate in junctional epidermolysis bullosa. J Cutan Pathol 2015; 42:559–63.

27. Nomura M, Hamasaki YI, Katayama I, Abe K, Niikawa N, Yoshiura KI. Eosinophil infiltration in three patients with generalized atrophic benign epidermolysis bullosa from a Japanese family: molecular genetic and immunohistochemical studies. J Hum Genet 2005; 50:483–9.

28. Roth RR, Smith KJ, James WD. Eosinophilic infiltrates in epidermolysis bullosa. Arch Dermatol 1990; 126:1191–4.

29. Franzke CW, Bruckner-Tuderman L, Blobel CP. Shedding of collagen XVII/BP180 in skin depends on both ADAM10 and ADAM9. J Biol Chem 2009; 284:23386–96.

30. Galiger C, Löffek S, Stemmler MP, Kroeger JK, Mittapalli VR, Fauth L, et al. Targeting of Cell Surface Proteolysis of Collagen XVII Impedes Squamous Cell Carcinoma Progression. Mol Ther 2018; 26:17–30.

31. Nishie W. Collagen XVII Processing and Blistering Skin Diseases. Acta Derm Venereol 2020; 100:adv00054.

32. Moilanen JM, Kokkonen N, Löffek S, Väyrynen JP, Syväniemi E, Hurskainen T, et al. Collagen XVII expression correlates with the invasion and metastasis of colorectal cancer. Hum Pathol 2015; 46:434–42.

33. Franzke CW, Tasanen K, Borradori L, Huotari V, Bruckner-Tuderman L. Shedding of collagen XVII/BP180: structural motifs influence cleavage from cell surface. J Biol Chem 2004; 279:24521–9.

34. Schumann H, Baetge J, Tasanen K, Wojnarowska F, Schäcke H, Zillikens D, et al. The shed ectodomain of collagen XVII/BP180 is targeted by autoantibodies in different blistering skin diseases. Am J Pathol 2000; 156:685–95.

35. Valdebran M, Kowalski EH, Kneiber D, Li J, Kim J, Doan L, et al. Epidermal expression of eotaxins and thymic stromal lymphopoietin in eosinophil rich dermatoses. Arch Dermatol Res 2019; 311:705–10.

36. Pigors M, Patzelt S, Reichhelm N, Dworschak J, Khil’chenko S, Emtenani S, et al. Bullous pemphigoid induced by IgG targeting type XVII collagen non-NC16A/NC15A extracellular domains is driven by Fc gamma receptor-and complement-mediated effector mechanisms and is ameliorated by neonatal Fc receptor blockade. J Pathol 2023.

37. Borradori L, Van Beek N, Feliciani C, Tedbirt B, Antiga E, Bergman R, et al. Updated S2 K guidelines for the management of bullous pemphigoid initiated by the European Academy of Dermatology and Venereology (EADV). J Eur Acad Dermatol Venereol 2022; 36:1689–704.

38. Hou B, Reizis B, DeFranco AL. Toll-like receptors activate innate and adaptive immunity by using dendritic cell-intrinsic and -extrinsic mechanisms. Immunity 2008; 29:272–82.

39. Swindell WR, Beamer MA, Sarkar MK, Loftus S, Fullmer J, Xing X, et al. RNA-Seq Analysis of IL-1B and IL-36 Responses in Epidermal Keratinocytes Identifies a Shared MyD88-Dependent Gene Signature. Front Immunol 2018; 9:80.

40. Dickson MA, Hahn WC, Ino Y, Ronfard V, Wu JY, Weinberg RA, et al. Human keratinocytes that express hTERT and also bypass a p16(INK4a)-enforced mechanism that limits life span become immortal yet retain normal growth and differentiation characteristics. Mol Cell Biol 2000; 20:1436–47.

41. Arnette C, Koetsier JL, Hoover P, Getsios S, Green KJ. In Vitro Model of the Epidermis: Connecting Protein Function to 3D Structure. Methods Enzymol 2016; 569:287–308.

42. Depianto D, Kerns ML, Dlugosz AA, Coulombe PA. Keratin 17 promotes epithelial proliferation and tumor growth by polarizing the immune response in skin. Nat Genet 2010; 42:910–4.

43. Hunyadi J, Simon M, Jr., Kenderessy AS, Dobozy A. Expression of monocyte/macrophage markers (CD13, CD14, CD68) on human keratinocytes in healthy and diseased skin. J Dermatol 1993; 20:341–5.

44. Schäbitz A, Hillig C, Mubarak M, Jargosch M, Farnoud A, Scala E, et al. Spatial transcriptomics landscape of lesions from non-communicable inflammatory skin diseases. Nat Commun 2022; 13:7729.

45. Bao L, Li J, Perez White BE, Patel PM, Amber KT. Inhibition of dipeptidyl-peptidase 4 induces upregulation of the late cornified envelope cluster in keratinocytes. Arch Dermatol Res 2022; 314:909–15.

46. Liu T, Wang Z, Xue X, Wang Z, Zhang Y, Mi Z, et al. Single-cell transcriptomics analysis of bullous pemphigoid unveils immune-stromal crosstalk in type 2 inflammatory disease. Nat Commun 2024; 15:5949.

47. Miyachi H, Wakabayashi S, Sugihira T, Aoyama R, Saijo S, Koguchi-Yoshioka H, et al. Keratinocyte IL-36 Receptor/MyD88 Signaling Mediates Malassezia-Induced IL-17-Dependent Skin Inflammation. J Infect Dis 2021; 223:1753–65.

48. Lee Y, Kim H, Kim S, Shin MH, Kim YK, Kim KH, et al. Myeloid differentiation factor 88 regulates basal and UV-induced expressions of IL-6 and MMP-1 in human epidermal keratinocytes. J Invest Dermatol 2009; 129:460–7.

49. Pivarcsi A, Bodai L, Réthi B, Kenderessy-Szabó A, Koreck A, Széll M, et al. Expression and function of Toll-like receptors 2 and 4 in human keratinocytes. Int Immunol 2003; 15:721–30.

50. Maglie R, Mercurio L, Morelli M, Madonna S, Salemme A, Baffa ME, et al. Interleukin-36 cytokines are overexpressed in the skin and sera of patients with bullous pemphigoid. Exp Dermatol 2023; 32:915–21.

51. Yamawaki Y, Kimura H, Hosoi T, Ozawa K. MyD88 plays a key role in LPS-induced Stat3 activation in the hypothalamus. Am J Physiol Regul Integr Comp Physiol 2010; 298:R403–10.

52. Nakagawa S, Matsumoto M, Katayama Y, Oguma R, Wakabayashi S, Nygaard T, et al. Staphylococcus aureus Virulent PSMα Peptides Induce Keratinocyte Alarmin Release to Orchestrate IL-17-Dependent Skin Inflammation. Cell Host Microbe 2017; 22:667–77.e5.

53. Kotnik N, Langner A, Meyer NH, Pas HH, Gibbs BF, Meijer JM, et al. Infiltration analysis of eosinophils and basophils and co-expression of CD69, CD63, IL-31 and IgE in patients with bullous and non-bullous pemphigoid. J Eur Acad Dermatol Venereol 2024; 38:e278–e81.

54. Lamberts A, Kotnik N, Meijer JM, van Kempen LC, Diercks GFH, Horváth B. Gene Expression Profiling Suggests that Complement Activation Is Important for Blister Formation in Bullous Pemphigoid. J Invest Dermatol 2023; 143:1591–4.e2.

55. Joly P, Roujeau JC, Benichou J, Picard C, Dreno B, Delaporte E, et al. A comparison of oral and topical corticosteroids in patients with bullous pemphigoid. N Engl J Med 2002; 346:321–7.

56. Jiang Y, Tsoi LC, Billi AC, Ward NL, Harms PW, Zeng C, et al. Cytokinocytes: the diverse contribution of keratinocytes to immune responses in skin. JCI Insight 2020; 5.

57. Kasprick A, Holtsche MM, Rose EL, Hussain S, Schmidt E, Petersen F, et al. The Anti-C1s Antibody TNT003 Prevents Complement Activation in the Skin Induced by Bullous Pemphigoid Autoantibodies. J Invest Dermatol 2018; 138:458–61.

58. Yan T, Xie Y, Liu Y, Shan Y, Wu X, Wang J, et al. Dupilumab effectively and rapidly treats bullous pemphigoid by inhibiting the activities of multiple cell types. Front Immunol 2023; 14:1194088.

59. Moghadam P, Tancrede E, Bouaziz JD, Kallout J, Bedane C, Begon E, et al. Efficacy and safety of dupilumab in bullous pemphigoid: a retrospective multicentric study of 36 patients. Br J Dermatol 2023; 189:244–6.

60. Learned C, Cohen SR, Cunningham K, Alsukait S, Santiago S, Lu J, et al. Long-term treatment outcomes and safety of dupilumab as a therapy for bullous pemphigoid: A multicenter retrospective review. J Am Acad Dermatol 2023; 89:378–82.

61. Abdat R, Waldman RA, de Bedout V, Czernik A, McLeod M, King B, et al. Dupilumab as a novel therapy for bullous pemphigoid: A multicenter case series. J Am Acad Dermatol 2020; 83:46–52.

62. Koga H, Teye K, Sugawara A, Tsutsumi M, Ishii N, Nakama T. Elevated levels of interleukin-9 in the serum of bullous pemphigoid: possible association with the pathogenicity of bullous pemphigoid. Front Immunol 2023; 14:1135002.

63. Kaplan MH, Hufford MM, Olson MR. The development and in vivo function of T helper 9 cells. Nat Rev Immunol 2015; 15:295–307.

64. Licona-Limón P, Arias-Rojas A, Olguín-Martínez E. IL-9 and Th9 in parasite immunity. Semin Immunopathol 2017; 39:29–38.

65. Yao W, Zhang Y, Jabeen R, Nguyen ET, Wilkes DS, Tepper RS, et al. Interleukin-9 is required for allergic airway inflammation mediated by the cytokine TSLP. Immunity 2013; 38:360–72.

66. Ma L, Xue HB, Guan XH, Shu CM, Zhang JH, Yu J. Possible pathogenic role of T helper type 9 cells and interleukin (IL)-9 in atopic dermatitis. Clin Exp Immunol 2014; 175:25–31.

67. Koch S, Sopel N, Finotto S. Th9 and other IL-9-producing cells in allergic asthma. Semin Immunopathol 2017; 39:55–68.

68. Licona-Limón P, Henao-Mejia J, Temann AU, Gagliani N, Licona-Limón I, Ishigame H, et al. Th9 Cells Drive Host Immunity against Gastrointestinal Worm Infection. Immunity 2013; 39:744–57.

69. Xue G, Jin G, Fang J, Lu Y. IL-4 together with IL-1β induces antitumor Th9 cell differentiation in the absence of TGF-β signaling. Nat Commun 2019; 10:1376

70. Soler D, Humphreys TL, Spinola SM, Campbell JJ. CCR4 versus CCR10 in human cutaneous TH lymphocyte trafficking. Blood 2003; 101:1677–82.

71. Sie C, Kant R, Peter C, Muschaweckh A, Pfaller M, Nirschl L, et al. IL-24 intrinsically regulates Th17 cell pathogenicity in mice. J Exp Med 2022; 219.

72. Li B, Jones LL, Geiger TL. IL-6 Promotes T Cell Proliferation and Expansion under Inflammatory Conditions in Association with Low-Level RORγt Expression. J Immunol 2018; 201:2934–46.

73. Mabrouk N, Tran T, Sam I, Pourmir I, Gruel N, Granier C, et al. CXCR6 expressing T cells: Functions and role in the control of tumors. Front Immunol 2022; 13:1022136.

74. Huffman AP, Lin JH, Kim SI, Byrne KT, Vonderheide RH. CCL5 mediates CD40-driven CD4+ T cell tumor infiltration and immunity. JCI Insight 2020; 5.

75. Yan Y, Chen R, Wang X, Hu K, Huang L, Lu M, et al. CCL19 and CCR7 Expression, Signaling Pathways, and Adjuvant Functions in Viral Infection and Prevention. Front Cell Dev Biol 2019; 7:212.

76. Herzog B, Pellet-Many C, Britton G, Hartzoulakis B, Zachary IC. VEGF binding to NRP1 is essential for VEGF stimulation of endothelial cell migration, complex formation between NRP1 and VEGFR2, and signaling via FAK Tyr407 phosphorylation. Mol Biol Cell 2011; 22:2766–76.

77. Malaponte G, Signorelli SS, Bevelacqua V, Polesel J, Taborelli M, Guarneri C, et al. Increased Levels of NF-kB-Dependent Markers in Cancer-Associated Deep Venous Thrombosis. PLoS One 2015; 10:e0132496.

78. Ferroni P, Palmirotta R, Riondino S, De Marchis ML, Nardecchia A, Formica V, et al. VEGF gene promoter polymorphisms and risk of VTE in chemotherapy-treated cancer patients. Thromb Haemost 2016; 115:143–51.

79. Schneeweiss MC, Merola JF, Wyss R, Silverberg JI, Mostaghimi A. Venous Thromboembolism in Patients With Bullous Pemphigoid. JAMA Dermatol 2023; 159:750–6.

80. Chen TL, Huang WT, Loh CH, Huang HK, Chi CC. Risk of Incident Venous Thromboembolism Among Patients With Bullous Pemphigoid or Pemphigus Vulgaris: A Nationwide Cohort Study With Meta-Analysis. J Am Heart Assoc 2023; 12:e029740.

81. Cugno M, Marzano AV, Bucciarelli P, Balice Y, Cianchini G, Quaglino P, et al. Increased risk of venous thromboembolism in patients with bullous pemphigoid. The INVENTEP (INcidence of VENous ThromboEmbolism in bullous Pemphigoid) study. Thromb Haemost 2016; 115:193–9.

82. Margaroli C, Bradley B, Thompson C, Brown MR, Giacalone VD, Bhatt L, et al. Distinct compartmentalization of immune cells and mediators characterizes bullous pemphigoid disease. Exp Dermatol 2020; 29:1191–8.

83. Ameglio F, D’Auria L, Cordiali-Fei P, Mussi A, Valenzano L, D’Agosto G, et al. Bullous pemphigoid and pemphigus vulgaris: correlated behaviour of serum VEGF, sE-selectin and TNF-alpha levels. J Biol Regul Homeost Agents 1997; 11:148–53.

84. Brown LF, Harrist TJ, Yeo KT, Ståhle-Bäckdahl M, Jackman RW, Berse B, et al. Increased expression of vascular permeability factor (vascular endothelial growth factor) in bullous pemphigoid, dermatitis herpetiformis, and erythema multiforme. J Invest Dermatol 1995; 104:744–9.

85. Jordan TJM, Chen J, Li N, Burette S, Wan L, Chen L, et al. The Eotaxin-1/CCR3 Axis and Matrix Metalloproteinase-9 Are Critical in Anti-NC16A IgE-Induced Bullous Pemphigoid. J Immunol 2023.

86. Dijkstra A, Postma DS, Noordhoek JA, Lodewijk ME, Kauffman HF, ten Hacken NH, et al. Expression of ADAMs ("a disintegrin and metalloprotease") in the human lung. Virchows Arch 2009; 454:441–9.

87. Naus S, Blanchet MR, Gossens K, Zaph C, Bartsch JW, McNagny KM, et al. The metalloprotease-disintegrin ADAM8 is essential for the development of experimental asthma. Am J Respir Crit Care Med 2010; 181:1318–28.

88. Chen J, Deng L, Dreymüller D, Jiang X, Long J, Duan Y, et al. A novel peptide ADAM8 inhibitor attenuates bronchial hyperresponsiveness and Th2 cytokine mediated inflammation of murine asthmatic models. Sci Rep 2016; 6:30451.

89. Matsuno O, Ono E, Ueno T, Takenaka R, Nishitake T, Hiroshige S, et al. Increased serum ADAM8 concentration in patients with drug-induced eosinophilic pneumonia-ADAM8 expression depends on a the allergen route of entry. Respir Med 2010; 104:34–9.

90. Higuchi Y, Yasui A, Matsuura K, Yamamoto S. CD156 transgenic mice. Different responses between inflammatory types. Pathobiology 2002; 70:47–54.

91. Schmidt E, Bröcker EB, Zillikens D. High levels of soluble CD23 in blister fluid of patients with bullous pemphigoid. Arch Dermatol 1995; 131:966–7.

92. Alam R, Stafford S, Forsythe P, Harrison R, Faubion D, Lett-Brown MA, et al. RANTES is a chemotactic and activating factor for human eosinophils. J Immunol 1993; 150:3442–8.

93. Frezzolini A, Teofoli P, Cianchini G, Barduagni S, Ruffelli M, Ferranti G, et al. Increased expression of eotaxin and its specific receptor CCR3 in bullous pemphigoid. Eur J Dermatol 2002; 12:27–31.

94. Bornscheuer E, Zillikens D, Schröder JM, Sticherling M. Lack of expression of interleukin 8 and RANTES in autoimmune bullous skin diseases. Dermatology 1999; 198:118–21.

95. Nakashima H, Fujimoto M, Asashima N, Watanabe R, Kuwano Y, Yazawa N, et al. Serum chemokine profile in patients with bullous pemphigoid. Br J Dermatol 2007; 156:454–9.

96. Patel PM, Jones VA, Kridin K, Amber KT. The role of Dipeptidyl Peptidase-4 in cutaneous disease. Exp Dermatol 2021; 30:304–18.

97. Chakievska L, Holtsche MM, Künstner A, Goletz S, Petersen BS, Thaci D, et al. IL-17A is functionally relevant and a potential therapeutic target in bullous pemphigoid. J Autoimmun 2019; 96:104–12.

98. Baggio LL, Varin EM, Koehler JA, Cao X, Lokhnygina Y, Stevens SR, et al. Plasma levels of DPP4 activity and sDPP4 are dissociated from inflammation in mice and humans. Nat Commun 2020; 11:3766.

99. Mentlein R. Dipeptidyl-peptidase IV (CD26)--role in the inactivation of regulatory peptides. Regul Pept 1999; 85:9–24.

100. Struyf S, Proost P, Schols D, De Clercq E, Opdenakker G, Lenaerts JP, et al. CD26/dipeptidyl-peptidase IV down-regulates the eosinophil chemotactic potency, but not the anti-HIV activity of human eotaxin by affecting its interaction with CC chemokine receptor 3. J Immunol 1999; 162:4903–9.

101. Nätynki A, Kokkonen N, Tuusa J, Ohlmeier S, Bergmann U, Tasanen K. Proteomic changes related to actin cytoskeleton function in the skin of vildagliptin-treated mice. J Dermatol Sci 2024; 113:121–9.

102. Goletz S, Probst C, Komorowski L, Radzimski C, Mindorf S, Holtsche MM, et al. Sensitive and specific assay for the serological diagnosis of anti-p200 pemphigoid based on the recombinant laminin β4 subunit. Br J Dermatol 2024.

103. Goletz S, Pigors M, Lari TR, Hammers CM, Wang Y, Emtenani S, et al. Laminin β4 is a constituent of the cutaneous basement membrane zone and additional autoantigen of anti-p200 pemphigoid. J Am Acad Dermatol 2024; 90:790–7.

104. Nishie W. Dipeptidyl peptidase IV inhibitor-associated bullous pemphigoid: a recently recognized autoimmune blistering disease with unique clinical, immunological and genetic characteristics. Immunol Med 2019; 42:22–8.

105. Ständer S, Hammers CM, Vorobyev A, Schmidt E, Zillikens D, Ghorbanalipoor S, et al. The impact of lesional inflammatory cellular infiltrate on the phenotype of bullous pemphigoid. J Eur Acad Dermatol Venereol 2021; 35:1702–11.

106. Guerrero-Juarez CF, Schilf P, Li J, Zappia MP, Bao L, Patel PM, et al. C-type lectin receptor expression is a hallmark of neutrophils infiltrating the skin in epidermolysis bullosa acquisita. Frontiers in Immunology 2023; 14.

