## Supplementary figures and images for "IgG autoantibodies in bullous pemphigoid directly induce a pathogenic MyD88-dependent pro-inflammatory response in keratinocytes"

### Supplementary Figure 1

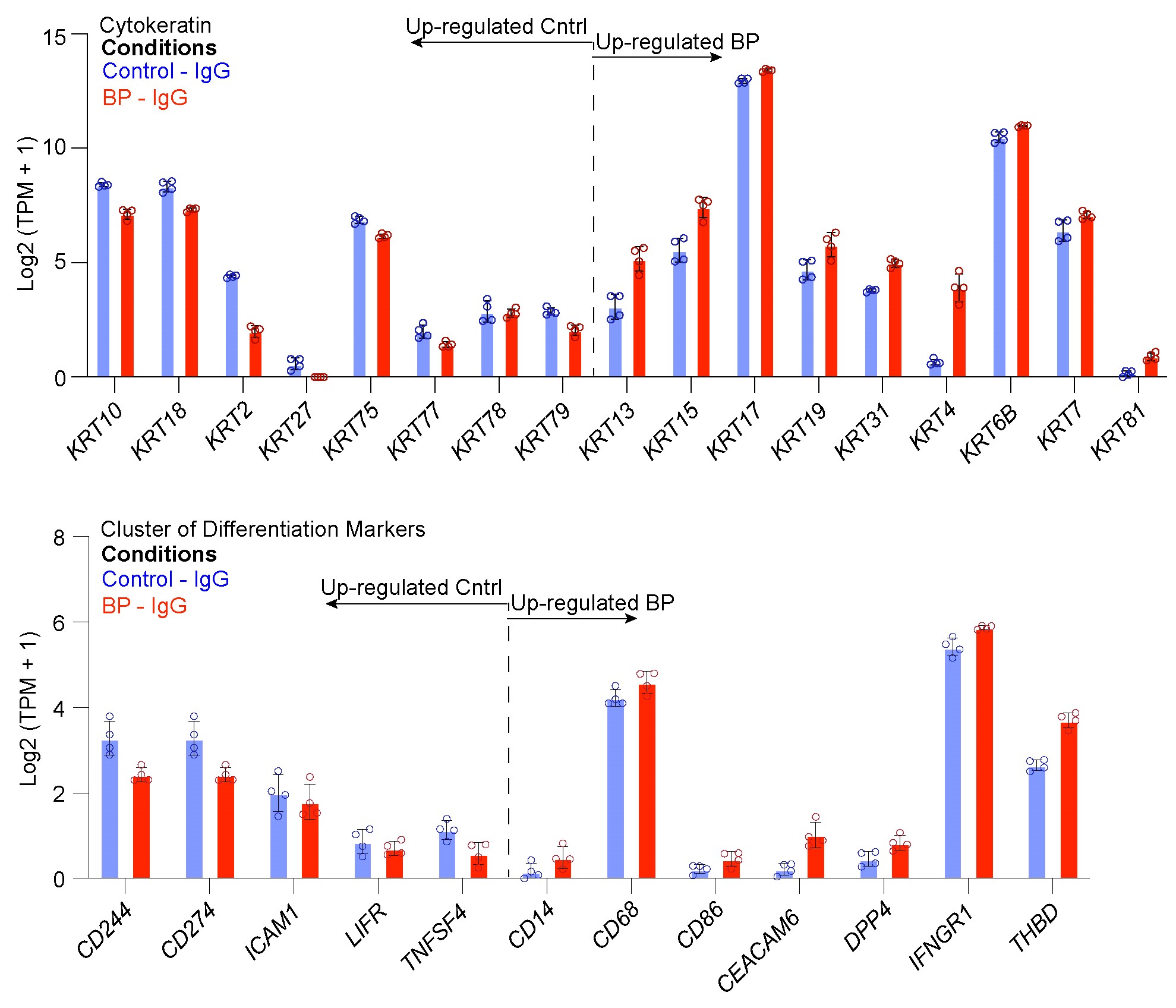

### Supplementary Figure 2

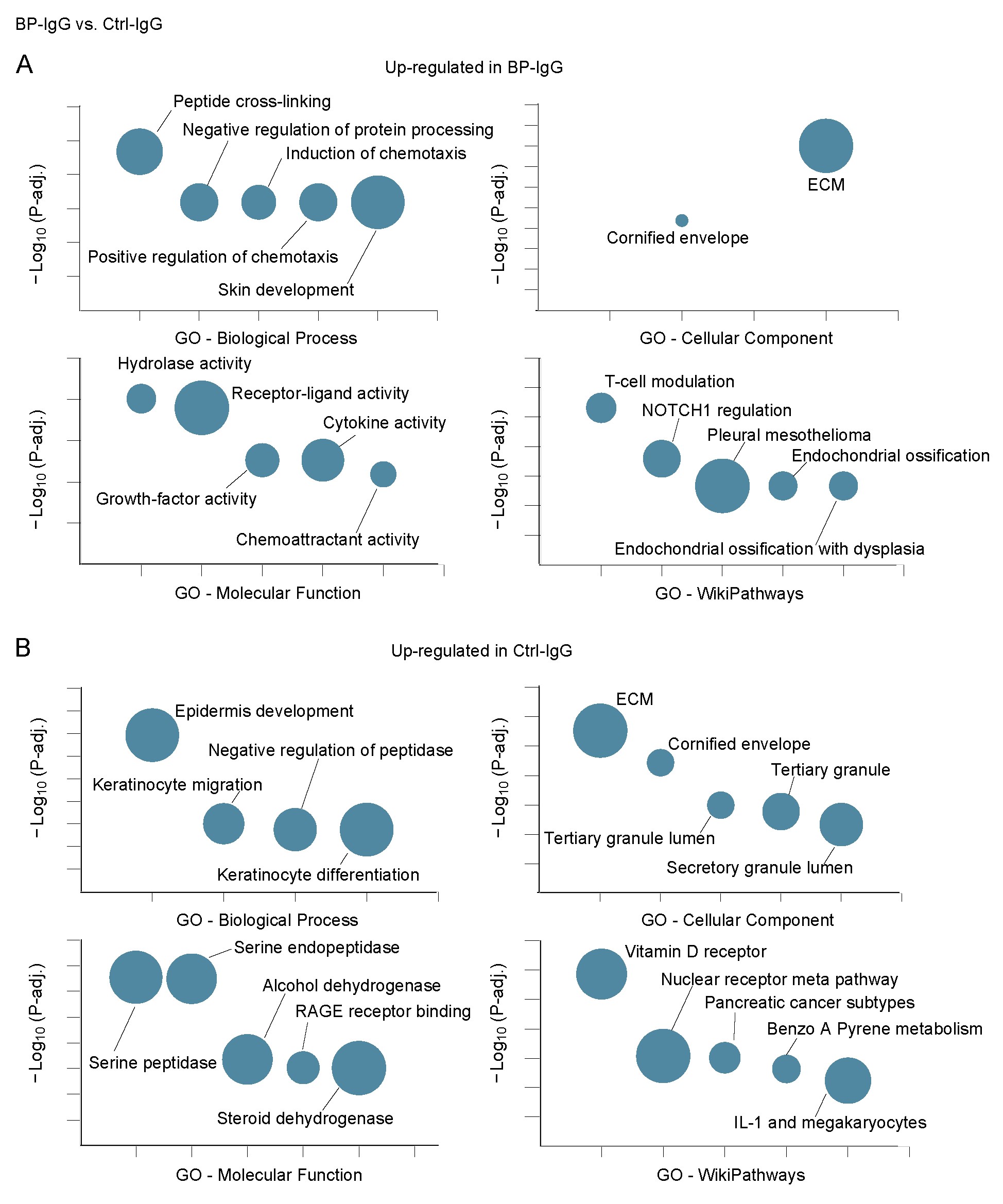

### Supplementary Figure 3

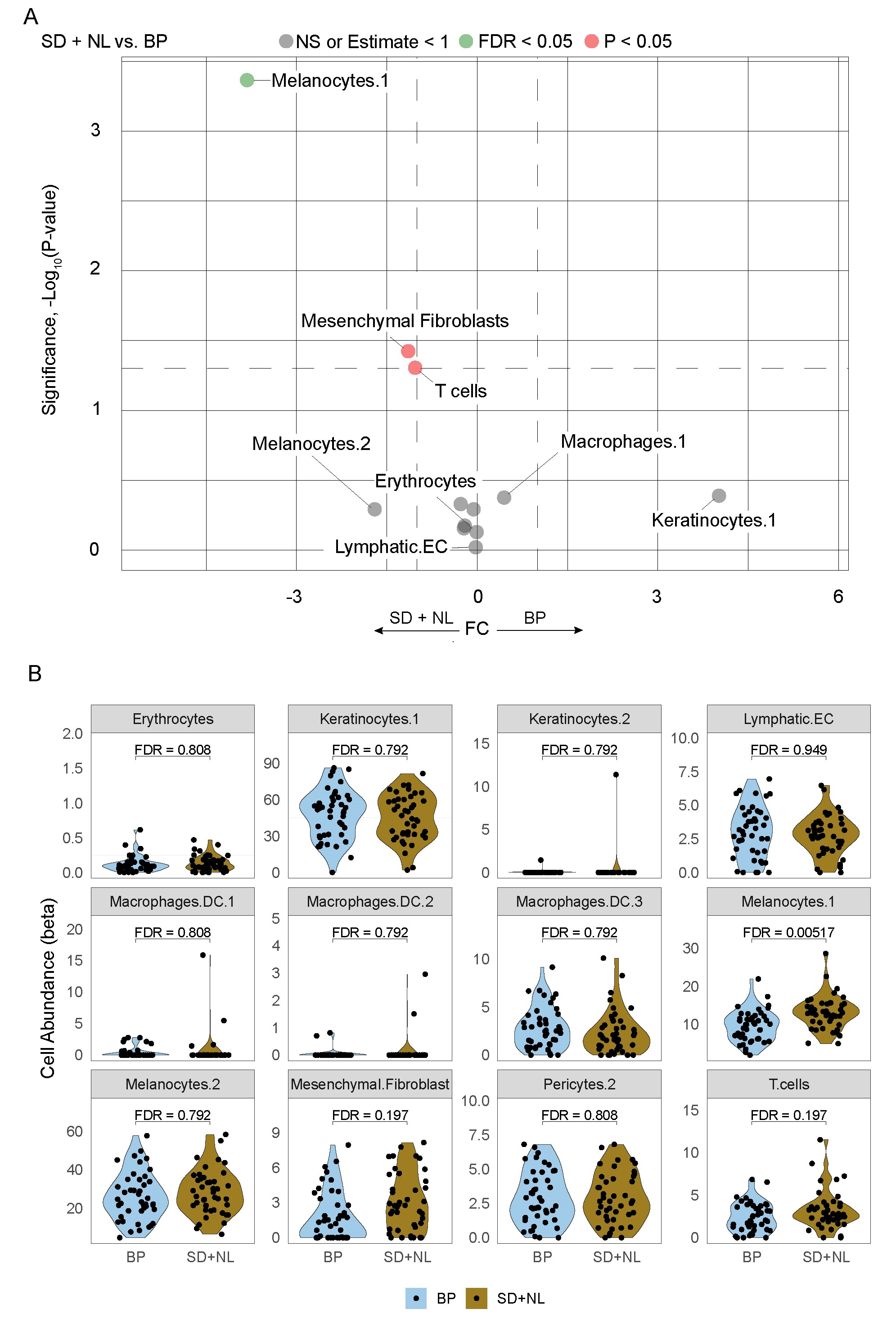

### Supplementary Figure 4

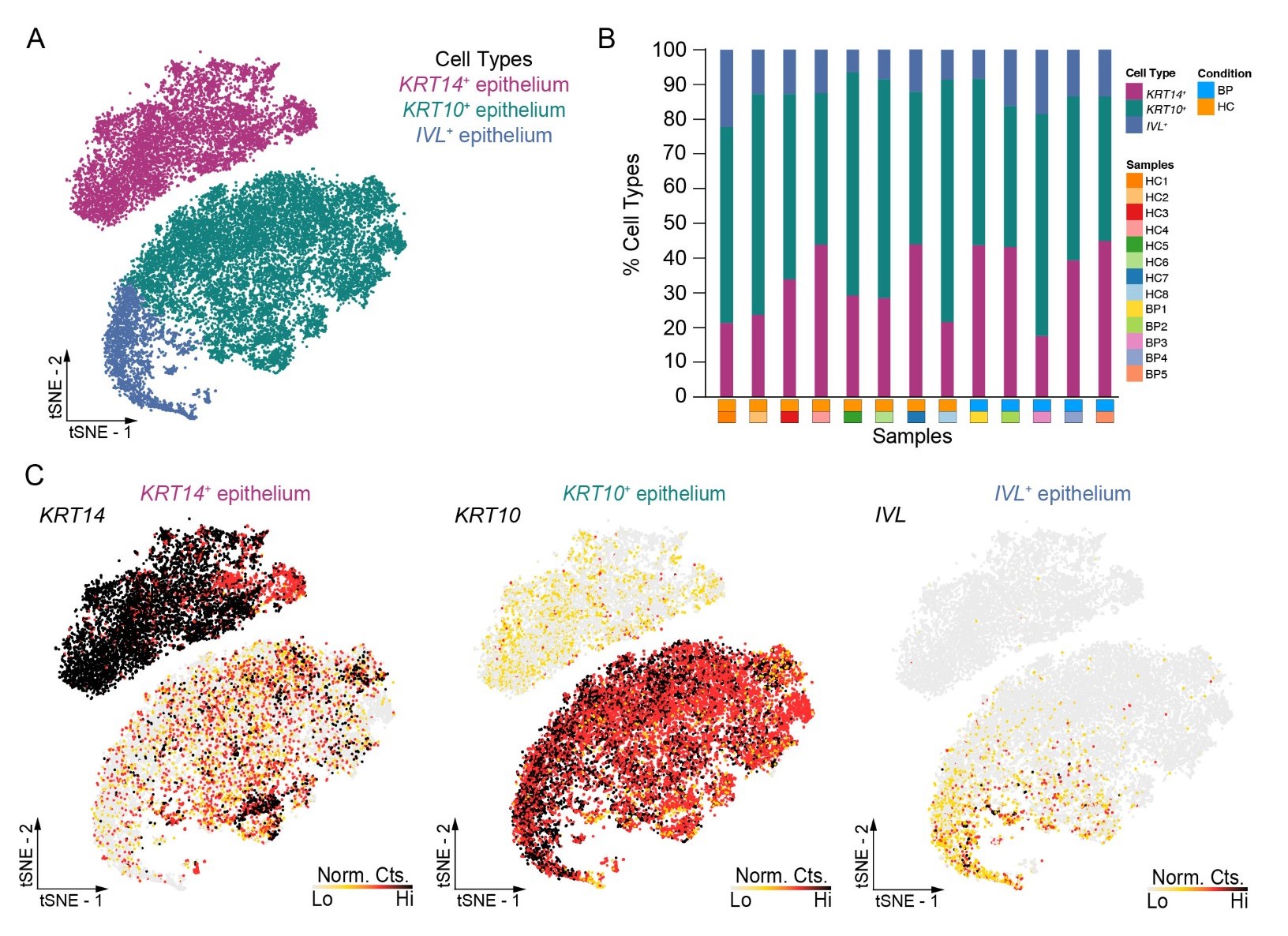

### Supplementary Figure 5

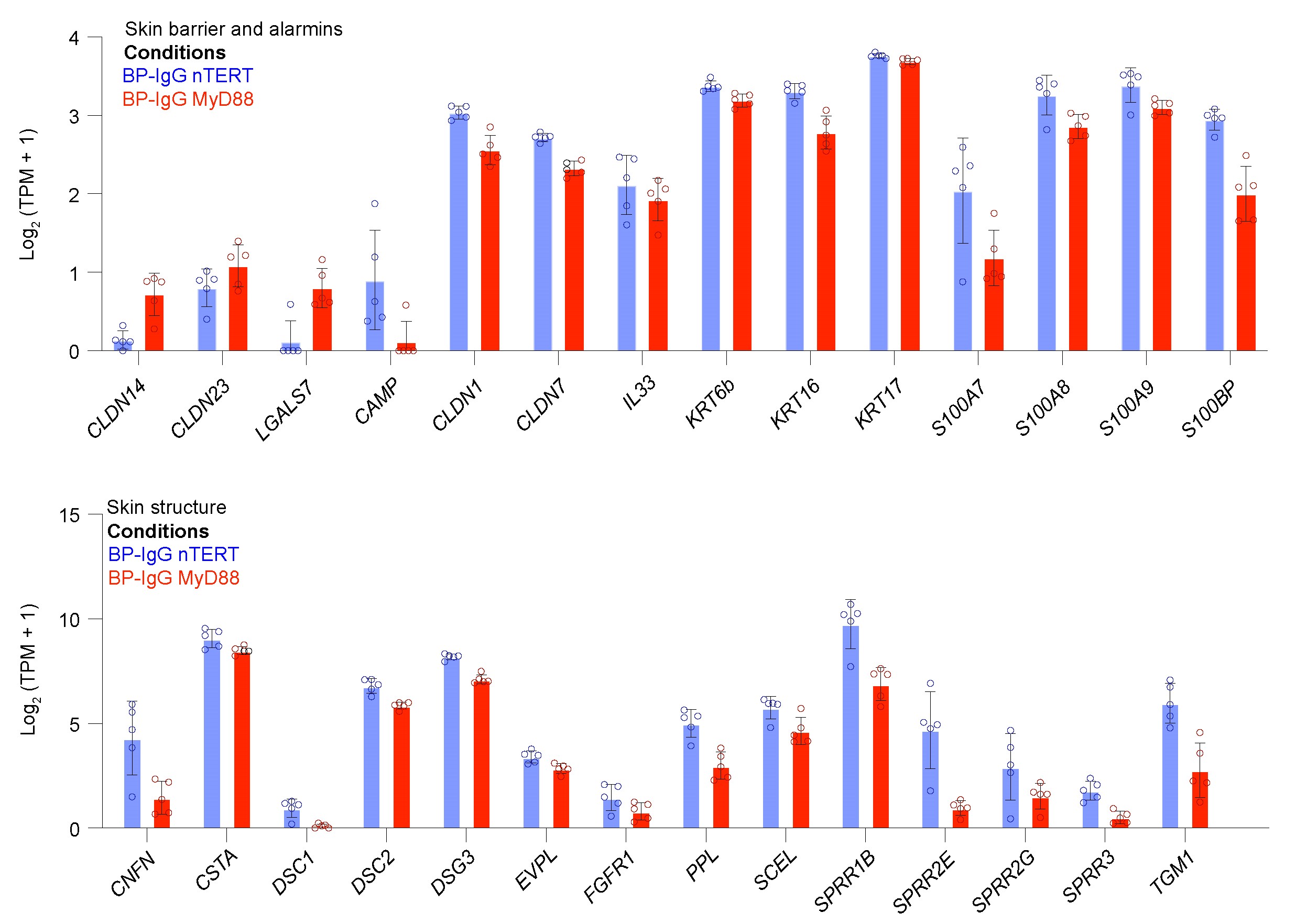

### Supplementary Figure 6

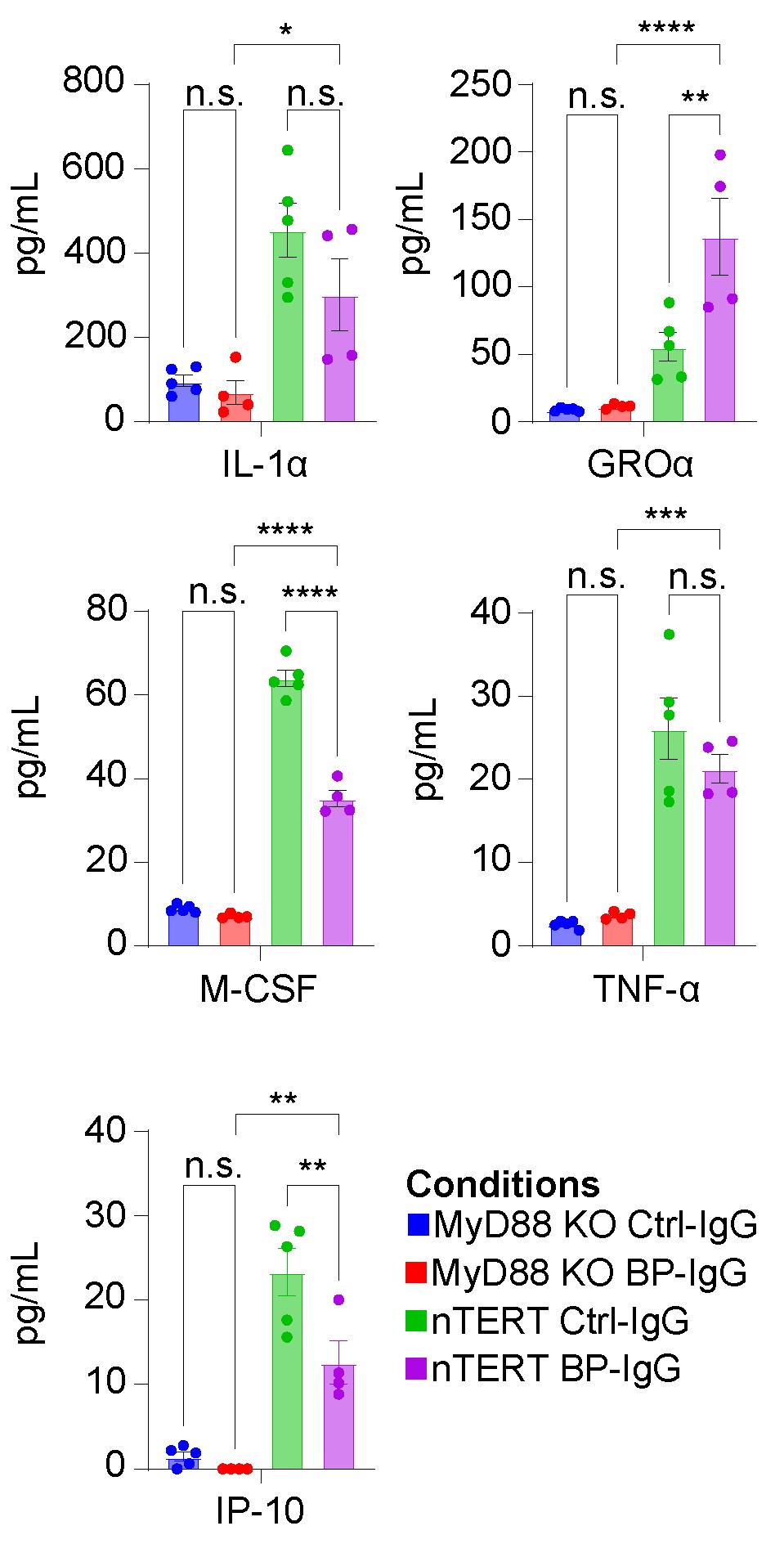
