## Supplementary Materials and Methods for "IgG autoantibodies in bullous pemphigoid directly induce a pathogenic MyD88-dependent pro-inflammatory response in keratinocytes"

**RNA Extraction:** RNA was extracted from keratinocytes using a miRNeasy Mini Kit according to the manufacturer’s instructions (Qiagen Inc., Germantown, MD, USA). On-Column DNase I Digestion was used to prevent genomic DNA contamination.

**RT-PCR:** Confirmation of MMP9 expression was performed as previously described^5^. After reverse transcription, primers targeting MMP9 (Forward primer: 3’-TCT TCC CTG GAG ACC TGA GA-5’ and reverse primer: 3’-ATT TCG ACT CTC CAC GCA TC-5’) were used for quantitative real-time PCR.

Primers were synthesized byIDT (Coralville, Iowa, USA). Data for qPCR were generated separate cohort of BP-IgG patients and keratinocytes.

**Bulk RNA-seq:** RNA quantity was determined using a Nanodrop 1000 spectrophotometer (Nanodrop, Wilmington, DE, USA). RNA integrity number (RIN) and concentration were determined using an Agilent 2100 Bioanalyzer (Agilent Technologies, Inc., Santa Clara, California, USA). A minimum RIN of 7 was required for further processing. Poly (A) mRNA was subsequently extracted and enriched from total RNA utilizing oligo (dT)-attached magnetic beads according to the manufacturer’s instructions (Illumina, San Diego, California, USA). Enriched and purified mRNA was fragmented into approximately 200 nt short mRNA, followed by first-strand cDNA synthesis using random hexamers as primers. Second-strand cDNA was constructed in a buffer containing dNTPs, DNA polymerase I, and RNase H. Appropriate fragments were isolated and enriched by PCR amplification. cDNA libraries were then sequenced using an Illumina NovaSeq sequencing platform with a read length of 2 x 150 and approximately 20 million reads per sample.

**Tissue Microarray Generation and Spatial Transcriptomics:** We utilized the Rush University Department of Pathology’s search feature to identify retrospective formalin fixed paraffin embedded (FFPE) tissue sections consistent with BP, spongiotic dermatitis with eosinophils (SD), or age matched normal skin (NL). BP samples were further confirmed by a positive serologic test. Individual sections from each specimen were stained with H&E per standard protocol to confirm the initial impression and suitability for the generation of a tissue microarray.

Regions of interest (ROIs) were pre-selected by the principal investigator (K.A.) and reviewed by a pathologist (V.M.) for histological verification. The array layout (4 columns x 7 rows) was defined following specific requirements to arrange donor cores in the center of a standard recipient block in an area no larger than 35 mm x 14 mm per GeoMx size requirements. An automated microarrayer, the TMA Master (3DHISTECH, Ltd., Budapest, Hungary) was used to remove and transfer cylindrical tissue cores from each donor block (cases & controls) and re-embed them into a recipient TMA paraffin block. The donor tissue core diameter was 2 mm and the distance between donor tissue cores was 0.7 mm. At least one core was chosen per patient, with additional cores taken for distinct regions such as the presence or absence of blistering. Three TMA blocks were built with samples randomized across the three blocks. Once construction was completed, the finalized TMA blocks were tempered overnight in a CO2 incubator at 37°C, facing down onto a clean glass slide to allow closure of any gap between the donor tissue cores and the surrounding paraffin wax. After tempering, the TMA blocks were left overnight to cool at room temperature before the glass slides were removed, and the blocks were ready for further use.

Representation of at least the epithelium of all samples was confirmed by cutting serial sections and performing H&E staining, followed by slide scanning utilizing the Aperio AT2 Slide scanner (Leica Biosystems, Wetzlar, Germany). Samples for transcriptomic analysis were cut to 5 microns. Samples were stained with antibodies targeting EPX (kindly provided by Dr. Elizabeth Jacob) directly conjugated to Alexa Fluor 594 (Lightning-Link, Abcam), and neutrophil elastase conjugated to Alexa Fluor 647 (MAB9167, Novus, Centenial, CO, USA) with DAPI counter stain to identify potential intraepidermal granulocytes. We then utilized the Nanostring GeoMx system (Nanostring, Seattle, Washington, USA) following the manufacturer’s recommend protocol to generate ROI and libraries. ROIs were manually assigned based on identification of epithelial morphology, excluding intraepithelial granulocytes. For BP samples, ROIs were defined as either blistered or adherent skin. Of 96 epithelial ROIs, 7 were removed due to low gene detection rate (<10%) or low alignment) (n=5 BP, n=1 SD, n=1 NL). Background signal was detected using negative control probes. Normalization was performed by the Quartile 3 approach. Data and figure generation were subsequently performed in R (4.1.0). Spatial deconvolution^6^ was performed to estimate the relative abundance of cell types in each ROI. Due to a lack of granulocytes in normal skin, we utilized two scRNA-seq data sets: Solé-Boldo et al^7^ and Danaher et al^6^.

**Bioinformatics:**

*Bulk RNA-seq analyses:* Paired-end reads were aligned to the human genome (Hg38/gencode.v42) with bowtie (version 1.2.3) and quantified using the RNA-seq by Expectation-Maximization algorithm (RSEM) (version 1.3.3) using standard parameters^8^. Samples were filtered and only protein-coding and lncRNA-coding genes expressed at minimum 1 TPM in at least one sample in all biological replicates were considered for downstream analyses. A pseudocount of 0.1 was added and samples were Log-transformed. Pair-wise comparisons were performed using edgeR as per developer’s suggestions^9^. Gene ontology pathway analyses was performed with (Enrichr)^10-12^. Downstream visualization including heatmaps (complexheatmap)^13, 14^, principal component analyses, and volcano plots (enhancedvolcano), were performed as suggested by developer with minor modifications. *Spatial transcriptomics:* GeoMx-NGS RNA expression analysis was performed with Geomx tools as previously described^15, 16^. Briefly, GeoMx raw count files, and metadata (DCC, PKC, and annotation files) were loaded in R Studio (version). Quality control, filtering and normalization was then performed as suggested by the developer with minor modifications. Q3 normalized data was used for differential gene expression analyses and performed as previously described^15, 16^. Downstream visualizations and statistical analysis including dimensionality reduction with UMAP and t-SNE, heatmaps, volcano plots, pathway analyses with Enrichr ^10-12^, and differential expression analyses with liner mixed effect models were performed as before and as suggested by developer. Raw sequencing files have been uploaded to the GEO server.

*Pseudo-bulk scRNA-seq analyses.* We utilized the recently published scRNA-seq database described by Liu *et al^17^*. Briefly, Cell ranger output files of 8 healthy controls (HC) and 5 BP patients were downloaded from Zenodo database (accession code: 10924853). Count matrices were pre-processed and doublets/multiplets were removed using Single-Cell Remover of Doublets (Scrublet)^18^ (version 0.2.1). The resultant AnnData object was exported as compressed AnnData H5AD file and were further read in Seurat for downstream low-quality cell pruning^19^. Seurat objects were merged, normalized, and highly variable genes (features) and scaling were performed using SCTransform^20^ as described before^21^. Basal, suprabasal, and granular keratinocytes were identified based on their bona fide expression of KRT14, KRT10, and IVL expression respectively. Differential gene expression in all keratinocytes between HC and BP patients was determined by DESeq2 with minor modifications^22^. Downstream visualization including heatmaps (complexheatmap), volcano plots (enhancedvolcano), and two-dimensional t-distributed Stochastic Neighbor Embedding (tSNE) were performed as suggested by developer with minor modifications.

**Experimental Pemphigoid Model:** The experimental BP model was generated as a previously described^23^. Briefly, recombinant NC14-1 cDNA was generated using gene synthesis (Eurofins MWG Operon, Ebersberg, Germany). After codon optimization for *E. coli*, cDNA was cloned into the pET24d-N expression vector where the NC14-1 protein was expressed with a terminal-His tag. Following expression in E*. coli* the recombinant protein was purified by affinity chromatography using TALON metal affinity resin (TaKaRa, San Jose, CA, USA). Anti-NC14-1 IgG was generated by immunization of New Zealand white rabbits (Kaneka Eurogentec S.A. Seraing, Belgium) with recombinant NC14-1. Total IgG from immunized rabbit sera was affinity purified using protein G Sepharose (Genscript, Piscataway, NJ, USA). Age- and sex-matched *Krt14*-Cre^+^;*Myd88^fl/fl^* or *Krt14*-Cre^-^;*Myd88^fl/fl^* littermates aged 8-12 weeks were injected intraperitoneally (i.p.) thrice weekly for 2 weeks to induce a cutaneous phenotype. The percentage of affected body surface area (ABSA) was measured by an investigator blinded to the experimental conditions on days 0, 4, 9, and 14. Disease scoring was modified from a previous mouse model of epidermolysis bullosa acquisita^24^. The percentage of ABSA was multiplied by 0.5 for alopecia, or 1.0 for erythema, crusting or erosions. Initial power analysis was performed to determine sizing using α= 0.05 and β= 0.8, with an anticipated mean disease surface area of 5 (±1) and an expected observed severity of 3.

| Species | Format | Analyte | Manufacturer/Catalog |
| --- | --- | --- | --- |
| Human | Luminex | sCD40L, EGF, Eotaxin, FGF-2, FLT-3 Ligand, Fractalkine, G-CSF, GM-CSF, GROα, IFN-α2, IFN-γ, IL-1α, IL-1β, IL-1RA, IL-2, IL-3, IL-4, IL-5, IL-6, IL-7, IL-8, IL-9, IL-10, IL-12(p40), IL-12(p70), IL-13, IL-15, IL-17A, IL-17E/IL-25, IL-17F, IL-18, IL-22, IL-27, IP-10, MCP-1, MCP-3, M-CSF, MDC, MIG/CXCL9, MIP-1α, MIP-1β, PDGF-AA, PDGF-AB/BB, RANTES, TGFα, TNF-α, TNF-β, and VEGF-A | MilliporeSigma,  HCYTA-60K-PX48 |
| Human | Luminex | 6CKine, APRIL, BAFF, BCA-1, CCL28, CTACK, CXCL16, ENA-78, Eotaxin-2, Eotaxin-3, GCP-2, Granzyme A, Granzyme B, HMGB1, I-309, I-TAC, IFNβ, IFNω, IL-11, IL-16, IL-20, IL-21, IL-23, IL-24, IL-28A, IL-29, IL-31, IL-33, IL-34, IL-35, LIF, Lymphotactin, MCP-2, MCP-4, MIP-1δ, MIP-3α, MIP-3β, MPIF-1, Perforin, sCD137, SCF, SDF-1, sFAS, sFASL, TARC, TPO, TRAIL, and TSLP | MilliporeSigma  HCYTB-60K-PXBK48 |
| Human | Luminex | MMP-1, MMP-2, MMP-3, MMP-7, MMP-8, MMP-9, MMP-10, MMP-12 and MMP-13 | R&D Systems  LMPM000 |
| Human | Luminex | TIMP-1, TIMP-2, TIMP-3 and TIMP-4 | R&D Systems  LKT003 |
| Human | Luminex | TGF-β1, TGF-β2, TGF-β3 | MilliporeSigma,  TGFBMAG-64K-03 |
| Human | ELISA | CD26 (EHDPP4) | Thermofisher  EHDPP4 |
| Human | ELISA | C1s (ELH-C1S-1) | RayBiotech  ELH-C1S-1 |
| Human | ELISA | IL-36G (EHIL36G) | Thermofisher  EHIL36G |
| Human | ELISA | ADAM8 (orb562354) | Bioorbyt  orb562354 |
| Mouse | Luminex | Eotaxin, G-CSF, GM-CSF, IFNγ, IL-1α, IL-1β, IL-2, IL-3, IL-4, IL-5, IL-6, IL-7, IL-9, IL-10, IL-12(p40), IL-12(p70), IL-13, IL-15, IL-17, IP-10, KC, LIF, LIX, MCP-1, M-CSF, MIG, MIP-1α, MIP-1β, MIP-2, RANTES, TNFα, and VEGF | MilliporeSigma  MCYTMAG-70K-PX32 |

Supplemental References:

1.	Hou B, Reizis B, DeFranco AL. Toll-like receptors activate innate and adaptive immunity by using dendritic cell-intrinsic and -extrinsic mechanisms. Immunity 2008; 29:272-82.

2.	Swindell WR, Beamer MA, Sarkar MK, Loftus S, Fullmer J, Xing X, et al. RNA-Seq Analysis of IL-1B and IL-36 Responses in Epidermal Keratinocytes Identifies a Shared MyD88-Dependent Gene Signature. Front Immunol 2018; 9:80.

3.	Dickson MA, Hahn WC, Ino Y, Ronfard V, Wu JY, Weinberg RA, et al. Human keratinocytes that express hTERT and also bypass a p16(INK4a)-enforced mechanism that limits life span become immortal yet retain normal growth and differentiation characteristics. Mol Cell Biol 2000; 20:1436-47.

4.	Arnette C, Koetsier JL, Hoover P, Getsios S, Green KJ. In Vitro Model of the Epidermis: Connecting Protein Function to 3D Structure. Methods Enzymol 2016; 569:287-308.

5.	Bao L, Li J, Solimani F, Didona D, Patel PM, Li X, et al. Subunit-Specific Reactivity of Autoantibodies Against Laminin-332 Reveals Direct Inflammatory Mechanisms on Keratinocytes. Front Immunol 2021; 12:775412.

6.	Danaher P, Kim Y, Nelson B, Griswold M, Yang Z, Piazza E, et al. Advances in mixed cell deconvolution enable quantification of cell types in spatial transcriptomic data. Nat Commun 2022; 13:385.

7.	Solé-Boldo L, Raddatz G, Schütz S, Mallm JP, Rippe K, Lonsdorf AS, et al. Single-cell transcriptomes of the human skin reveal age-related loss of fibroblast priming. Commun Biol 2020; 3:188.

8.	Li B, Dewey CN. RSEM: accurate transcript quantification from RNA-Seq data with or without a reference genome. BMC Bioinformatics 2011; 12:323.

9.	Robinson MD, McCarthy DJ, Smyth GK. edgeR: a Bioconductor package for differential expression analysis of digital gene expression data. Bioinformatics 2010; 26:139-40.

10.	Kuleshov MV, Jones MR, Rouillard AD, Fernandez NF, Duan Q, Wang Z, et al. Enrichr: a comprehensive gene set enrichment analysis web server 2016 update. Nucleic Acids Res 2016; 44:W90-7.

11.	Xie Z, Bailey A, Kuleshov MV, Clarke DJB, Evangelista JE, Jenkins SL, et al. Gene Set Knowledge Discovery with Enrichr. Curr Protoc 2021; 1:e90.

12.	Chen EY, Tan CM, Kou Y, Duan Q, Wang Z, Meirelles GV, et al. Enrichr: interactive and collaborative HTML5 gene list enrichment analysis tool. BMC Bioinformatics 2013; 14:128.

13.	Gu Z, Eils R, Schlesner M. Complex heatmaps reveal patterns and correlations in multidimensional genomic data. Bioinformatics 2016; 32:2847-9.

14.	Gu Z. Complex heatmap visualization. Imeta 2022; 1:e43.

15.	Zimmerman SM, Fropf R, Kulasekara BR, Griswold M, Appelbe O, Bahrami A, et al. Spatially resolved whole transcriptome profiling in human and mouse tissue using Digital Spatial Profiling. Genome Res 2022; 32:1892-905.

16.	Merritt CR, Ong GT, Church SE, Barker K, Danaher P, Geiss G, et al. Multiplex digital spatial profiling of proteins and RNA in fixed tissue. Nat Biotechnol 2020; 38:586-99.

17.	Liu T, Wang Z, Xue X, Wang Z, Zhang Y, Mi Z, et al. Single-cell transcriptomics analysis of bullous pemphigoid unveils immune-stromal crosstalk in type 2 inflammatory disease. Nat Commun 2024; 15:5949.

18.	Wolock SL, Lopez R, Klein AM. Scrublet: Computational Identification of Cell Doublets in Single-Cell Transcriptomic Data. Cell Syst 2019; 8:281-91.e9.

19.	Fan J, Salathia N, Liu R, Kaeser GE, Yung YC, Herman JL, et al. Characterizing transcriptional heterogeneity through pathway and gene set overdispersion analysis. Nat Methods 2016; 13:241-4.

20.	Hafemeister C, Satija R. Normalization and variance stabilization of single-cell RNA-seq data using regularized negative binomial regression. Genome Biol 2019; 20:296.

21.	Guerrero-Juarez CF, Schilf P, Li J, Zappia MP, Bao L, Patel PM, et al. C-type lectin receptor expression is a hallmark of neutrophils infiltrating the skin in epidermolysis bullosa acquisita. Frontiers in Immunology 2023; 14.

22.	Love MI, Huber W, Anders S. Moderated estimation of fold change and dispersion for RNA-seq data with DESeq2. Genome Biol 2014; 15:550.

23.	Pigors M, Patzelt S, Reichhelm N, Dworschak J, Khil'chenko S, Emtenani S, et al. Bullous pemphigoid induced by IgG targeting type XVII collagen non-NC16A/NC15A extracellular domains is driven by Fc gamma receptor- and complement-mediated effector mechanisms and is ameliorated by neonatal Fc receptor blockade. J Pathol 2023.

24.	Kasprick A, Bieber K, Ludwig RJ. Drug Discovery for Pemphigoid Diseases. Curr Protoc Pharmacol 2019; 84:e55.
