## Supplementary Tables and Data Legends for "IgG autoantibodies in bullous pemphigoid directly induce a pathogenic MyD88-dependent pro-inflammatory response in keratinocytes"

**sTable 1. Serotypes of included BP patients.**

|  | Age | DPP4i | BP180 | BP230 | IIF SSS |
| --- | --- | --- | --- | --- | --- |
| BP1 | 89 | No | * | * | * |
| BP2 | 80 | No | 126 | 115 | 40,960 |
| BP3 | 80 | No | 100 | 1 | 20 |
| BP4 | 80 | No | 24 | 36 | 5,120 |
| BP5 | 85 | No | Neg | 70 | 20,480 |
| BP6 | 85 | Yes | 59 | 0 | - |
| BP7 | 74 | No | 18 | 15 | 1,160 |
| BP8 | 85 | No | 78 | 62 | 20,480 |
| BP9 | 85 | No | 82 | 54 | * |
| BP10 | 64 | No | 93 | 4 | 2,560 |
| BP11 | 71 | No | Neg | 86 | +** |
| BP12 | 75 | No | Neg | Neg | +** |
| BP13 | 61 | No | Neg | 18 | +** |
| BP14 | 72 | No | Neg | Neg | +** |
| BP15 | 53 | No | 150 | Neg | +** |
| BP16 | 93 | No | 70 | 68 | +** |

*Prior serologies confirmatory of BP but not repeated at time of flare. ** Positive at 1:40 or greater on epidermal side of salt split human skin. BP – Bullous Pemphigoid, DPP4i – Dipeptidyl peptidase 4 inhibitor, IIF – Indirect immunofluorescence, SSS – Salt Split Skin.

**sTable 2. Patient demographics for histologic tissue used in tissue microarrays.**

| **Age** | **Gender** | **Location** | **DIF** |
| --- | --- | --- | --- |
| **Bullous Pemphigoid** | | | |
| 60 | M | Thigh | Linear C3 at DEJ, Focal IgG |
| 70 | F | Upper arm | Linear C3 and IgG (weak) |
| 83 | M | Forearm | Linear C3 and IgG |
| 78 | M | Arm | Linear C3 and IgG |
| 56 | M | Wrist | Linear C3 and IgG |
| 70 | F | Arm | Linear C3 and IgG |
| 66 | F | Arm | Linear IgG |
| 77 | M | Forearm | Linear C3 and IgG |
| 79 | F | Thigh | Linear C3 and IgG |
| 74 | M | Thigh | Linear C3 and IgG |
| 39 | M | Forearm | Linear C3 and IgG |
| 67 | M | Upper arm | Linear C3 and IgG |
| 71 | M | Upper arm | Linear C3 and IgG |
| 73 | M | Frontal scalp | Linear C3, IgG, IgM, and fibrinogen |
| 79 | M | Medial mid back | Linear C3 and IgG |
| 88 | F | Right hip | Linear C3 and IgG |
| 86 | F | Upper arm | Linear C3 and IgG |
| 82 | M | Posterior calf | Linear C3 |
| 97 | M | Arm | Linear C3 and IgG |
| 76 | M | Anterior arm | Linear C3 and IgG |
| **Spongiotic dermatitis with eosinophils** | | | |
| 67 | F | Lateral lower leg | **-** |
| 61 | F | Left neck | **-** |
| 65 | M | Thigh | Negative DIF |
| 48 | F | Upper thigh | **-** |
| 65 | F | Thigh | **-** |
| 71 | M | Abdomen/Thigh | **-** |
| 37 | M | Buttock | **-** |
| 54 | F | Posterior thigh | **-** |
| 59 | F | Arm | **-** |
| 25 | F | Upper back | Negative DIF |
| 23 | F | Breast | **-** |
| 44 | F | Upper arm | **-** |
| 70 | M | Abdomen | Negative DIF |
| 40 | F | Breast | **-** |
| 77 | F | Flank | Negative DIF |
| 28 | F | Hip | **-** |
| 83 | F | Hip | **-** |
| 74 | F | Arm | **-** |
| 55 | M | Back | **-** |
| 44 | M | Axilla | **-** |
| **Normal elderly skin** | | | |
| 82 | F | Mid back | **-** |
| 80 | M | Forehead | **-** |
| 73 | M | Forearm | **-** |
| 71 | M | Upper back | **-** |
| 77 | M | Central back | **-** |
| 72 | F | Central upper back | **-** |
| 71 | M | Forearm | **-** |
| 71 | M | Chest | **-** |
| 74 | M | Chest | **-** |
| 73 | M | Upper chest | **-** |
| 70 | M | Upper abdomen | **-** |
| 78 | M | Upper abdomen | **-** |
| 71 | F | Upper chest | **-** |
| 73 | F | Medial chest | **-** |
| 74 | M | Dorsal forearm | **-** |
| 75 | M | Forearm | **-** |
| 76 | M | Flank | **-** |
| 73 | F | Arm | **-** |
| 80 | F | Back | **-** |
| 88 | F | Temple | **-** |
| 82 | F | Leg | **-** |

*DEJ – Dermal epidermal junction, DIF – Direct immunofluorescence*

**Supplementary Data Description:**

**Supplemental data 1:** Proteins from supernatants of BP-IgG- vs control-IgG-treated primary keratinocytes meeting minimum detection thresholds. Data are shown as pg/ml.

**Supplementary data 2:** Significantly differentially expressed genes in BP- vs control IgG-treated primary human keratinocytes. Data shown are for changes greater than Log_2_F.C.(0.5) and P_adj_ < 0.05.

**Supplemental data 3:** Differential gene expression between detached vs attached epithelium in BP. Data shown are P_adj_ < 0.05.

**Supplemental data 4:** Overlapping gene profiles of BP- vs. control-IgG treated primary human keratinocytes with detached vs. attached epithelia in patients with BP.

**Supplemental data 5:** Differential gene expression of epithelia from attached BP skin and normal skin by spatial transcriptomics.

**Supplemental Data 6:** Overlap of differentially expressed genes in BP vs. normal skin. And BP-IgG vs. Control-IgG treated keratinocytes

**Supplemental data 7:** Differential gene expression between all BP vs. healthy control keratinocytes.

**Supplemental data 8:** Differential gene expression between of bioinformatically gated basal keratinocytes in BP vs. healthy control skin.

**Supplemental data 9:** Significantly differentially expressed genes in BP-IgG-treated MyD88 KO nTERT cells versus control cells. Data shown have Log_2_F.C. > 0.5 and P_adj_ <0.05 representative of n=4-5 per cohort.

**Supplemental data 10:** Significantly differentially expressed genes in BP-IgG- versus control-IgG-treated MyD88 KO nTERT cells. Data shown have Log_2_F.C. > 0.5 and P_adj_ <0.05 representative of n=4-5 per cohort.

**Supplemental data 11:** Supernatant protein levels in BP-IgG- or control-IgG-treated MyD88-deficient or wild type nTERT keratinocytes.
